# Tailoring Precise Genomic Integration Toward Isolate-to-Industry Strain Development for Scalable High-Titer Production of Polyhydroxyalkanoate

**DOI:** 10.64898/2026.07.23.735987

**Authors:** Peng Liu, Xin-Ying Xie, Yi-Hao Deng, Ze-Feng Li, Chi Wang, Hao Yang, Yu-Xi Li, Ling-Ling Zhao, Wei Situ, Hong-Wei Shen, Liu-Song Yu, Jin-Yan Lv, Yan-Chun Xiao, Yi-Na Lin, Jian-Wen Ye

## Abstract

Halophilic chassis has emerged as a promising biomanufacturing platform for industrial polyhydroxyalkanoate (PHA) production. However, challenges still remain in improving the production capacity, scalability and robustness, thereby lowering cost to meet market demands. Here, a high-performing halophilic strain *Halomonas* LY03 was isolated with over 38% glucose-to-PHA conversion rate and broad non-grain substrate utilization capability. Multidimensional tools, including algorithm-guided high-expression neutral integration site (HENIS) screening toolkit designated ‘SiteSeek’, stop codon (TAA)-dependent enhancement of gene expression and recombinase-mediated large-fragment (> 9 kb) genomic integration, were then developed to enable precise, efficient and interference-free genomic integrative expression. Using these tools, various chromosomally engineered strains were rapidly constructed to achieve high-level production of poly-3-hydroxybutyrate (PHB, 151 g L⁻¹) and poly(3-hydroxybutyrate-*co*-4-hydroxybutyrate) (P34HB, 139 g L⁻¹) under high cell-density fermentation (up to 186 g L^−1^ cell dry weight) in a 5-L bioreactor. Scalability was demonstrated at 2-m³ and 20-m³ industry-scale fermentations, yielding up to 134 g L⁻¹ PHB and 127 g L⁻¹ P34HB (6.1 mol% 4HB). Building on the proven robustness, a two-stage continuous fermentation (TCF) process was developed using a twin-bioreactor system at 5-L and 20-m³ scales, where stable and sustained PHA production lasted over 260 h and 160 h, respectively. Techno-economic analysis revealed a substantial cost-reduction space of 48% compared with conventional fed-batch process. This study demonstrates a successful paradigm for engineering a newly isolated strain toward robust, high-titer and cost-competitive PHA production across lab-to-industry scales.

## Introduction

To address mounting eco-environmental concerns arising from the widespread use of petrochemical plastics, particularly the human-health risks posed by microplastics pollution, the industrial biomanufacturing of polyhydroxyalkanoates (PHAs), one of the most promising alternatives capable of full-environment degradability, produced by engineered microbes using bio-based feedstocks such as sugars has emerged as a highly attractive and sustainable solution^1, 2^. To date, PHAs have been studied and used in versatile areas, such as packaging, 3D-printing, paper coating, biomedical implants, agricultural films and so on^3, 4^, due to the tunable and designable material properties powered by genetically reprogrammed microbial chassis like *Escherichia coli* (*E. coli*)^5^, *Cupriavidus necator* (*C. necator*)^6, 7^, the *Halomonas*^8^, etc^9^. For PHAs production using traditional chassis (e.g., *E. coli*, *C. necator*), the intractable risk of contamination and heavy energy consumption of sterilization represent long-standing but unsolved bottlenecks to cost reduction. In contrast, different halophiles, especially *Halomonas* strains, able to grow on saline and alkaline condition have been successfully isolated and engineered to produce diverse PHAs under unsterilized and open condition with reduced production cost^10, 11, 12, 13^.

Even with these improvements, the production cost of PHAs still remains stubbornly high for large-scale commercialization. Therefore, many efforts focusing on metabolic engineering of *Halomonas* chassis have been made to further improve the cost-competitiveness of PHAs. For instance, engineering larger cell size for PHA accumulation^14, 15^, manipulating the PHA granule size for simplified downstream extraction^16^, rebalancing the NADH/NAD^+^ ratio for enhanced PHA synthesis^17^, increasing the oxygen availability for improved PHA synthesis even under low-oxygen condition^18^, designing protein phase separation for enhanced glucose-to-PHA conversion^19^, engineering detoxification process for enhanced PHA copolymer from high concentration volatile fatty acid^20^, and so on. Meanwhile, isolating novel-type halophiles to produce PHAs using various non-grain substrates, such as food-derived wastes^21, 22, 23^, lignocellulose^24, 25, 26, 27, 28^, etc.^29, 30, 31, 32^, is also a promising strategy to develop a greener process of improved sustainability. Additionally, tailoring PHA synthesis (e.g., monomer composition, molecular weight, polydispersity index) to achieve a favorable trade-off between production cost and material properties, which is critical for enabling broad market adoption, is of great significance for realizing the end-to-end commercialization and full market application closure of PHAs. The flexible copolymer consisting of 3-hydroxybutyrate (3HB) and 4-hydroxybutyrate (4HB), namely P34HB, is recognized as a typical representative^33^. In recent years, advances in the metabolic engineering of P34HB production based on microbial chassis, especially for *Halomonas*, have substantially driven progresses in industrial PHA biomanufacturing (MedPHA Bioscience Co., Ltd., PhaBuilder Biotechnology Co., Ltd., CJ Biomaterials, Inc.).

Since the uncoordinated expression of 4HB synthesis pathway would generally lead to metabolic perturbation, even severe cytotoxicity^34^, the rationales of gene-expression level maintenance across the plasmid-to-genome system and efficiency of genomic integration generally determine the stability of P34HB-producing performance and scale-up robustness of engineered strains. However, these challenges remain unresolved despite extensive researches focusing on advancing genetic editing^35, 36, 37, 38^ and metabolic engineering tools and strategies^39, 40^ that have greatly accelerated strain development for *Halomonas*-based P34HB production^41, 42^. Nevertheless, almost no reported study can achieve high-level P34HB production with a titer over 100 g L^−1 41, 42^ under fed-batch condition except the brittle one PHB reached approximately 120 g L^−1 14^, which is still lower than that based on model chassis like *E. coli*^5^ and *C. necator*^6^.

Firstly, most developed genomic integration approaches for *Halomonas* strains rely on homologous recombination^43^, including CRISPR/Cas-mediated editing systems^44^. Their integration efficiency varies drastically across different host strains, and such methods generally perform poorly in non-model microorganisms^43, 45, 46^. The site-specific recombinase-directed integration has emerged as a powerful alternative of high-efficiency, host-independence and marker-free demand^47, 48, 49^, which has been broadly validated in various model and non-model strains^50, 51^, albeit without in halophilic species. Secondly, the ‘position effect’, which refers to dramatic variations in gene expression among distinct genomic integration loci, has been well documented in both model chassis such as *E. coli*^52^ and *yeast*^53, 54, 55, 56^, as well as the non-model strain *Halomonas* TD^44^. Nevertheless, useful strategies for identifying high-expression neutral integration sites (HENIS) are still urgently needed. Finally, as intracellular biomacromolecules, PHAs are almost exclusively produced via discontinuous fed-batch fermentation. This mode leaves little room for further boosting productivity and cutting operational costs. In contrast, developing continuous fermentation for PHA production can deliver higher space-time efficiency by eliminating downtime between batches, while also facilitating stable quality control of the final products^57, 58^.

In this work, we isolated a high-yield PHB-producing chassis strain, *Halomonas* LY03, and constructed an industrially applicable platform for PHA biomanufacturing. We also developed multidimensional genomic integration tools to enable rapid and precise chromosomal engineering of diverse industrial strains. These strains support antibiotic-free, high-titer and stable production of PHB and P34HB using glucose or glucose-xylose mixtures, which perform well across laboratory and industrial fermentation scales. More importantly, we established a two-stage continuous fermentation process based on a twin-bioreactor system. Compared with conventional fed-batch fermentation, this strategy enables long-term stable synthesis of PHB and P34HB, while greatly enhancing volumetric production efficiency and lowering production costs.

## Results

### Isolation and characterization of a high-performing halophilic chassis for PHA production

A customized six-step workflow was designed to isolate PHA-producing chassis in the saline-alkaline mud sampled from the salt lakes in Xinjiang, China (Fig. 1a). Guided by this strain screening pipeline, a total of 216 colonies with a diameter over 1 mm and low light transmittance (a typical phenotypic characteristic of intracellular PHA accumulation) were preliminarily isolated from massive microbial populations. Followed by growth assays in a 96-deep-well plate, 61 strains that satisfied the screening thresholds (OD_600_ > 1.7, 10% lower than positive control *Halomonas* TD01^9^ grown on the same condition; maximum specific growth rate (μₘₐₓ, h^−1^) > 0.35) were subjected to subsequent semi-quantification of PHA via flow cytometry analysis using a BODIPY (493/503) fluorescent staining method reported in previous studies^59, 60^. 21 Candidate strains with a fluorescence intensity (FI) higher than 12,000 (surpassing that of TD01) were thereby screened out. Combined with shake-flask fermentation test and duplicates removal by 16S rRNA sequencing (Figs. S1-S2), a dominant strain with the optimal PHB synthesis capacity, designated *Halomonas* LY03 (Table S3), was ultimately screened and identified. Further cultural medium optimization including glucose (carbon source) and urea (nitrogen source) supplementation dosages dramatically elevated PHB production and substrate conversion efficiency. In shake flask study, the optimized PHB titer reached 13.4 g L^−1^, with a glucose-to-PHB conversion efficiency up to 38.2% (Fig. 1b, Fig. S3). Moreover, LY03 exhibited prominent robustness in cell growth and PHB accumulation across diverse extreme or adverse cultural conditions (e.g., high pH, poor organic nutrition), and carbon feedstocks (Fig. 1c, Figs. S3-S4). Notably, the superior pH tolerance, reaching to 11 (Fig. S3f), enables the potential implementation of open, non-sterile fermentation under lower salt concentration, which substantially cuts down subsequent salt-containing wastewater disposal costs^61^.

**Figure 1.**
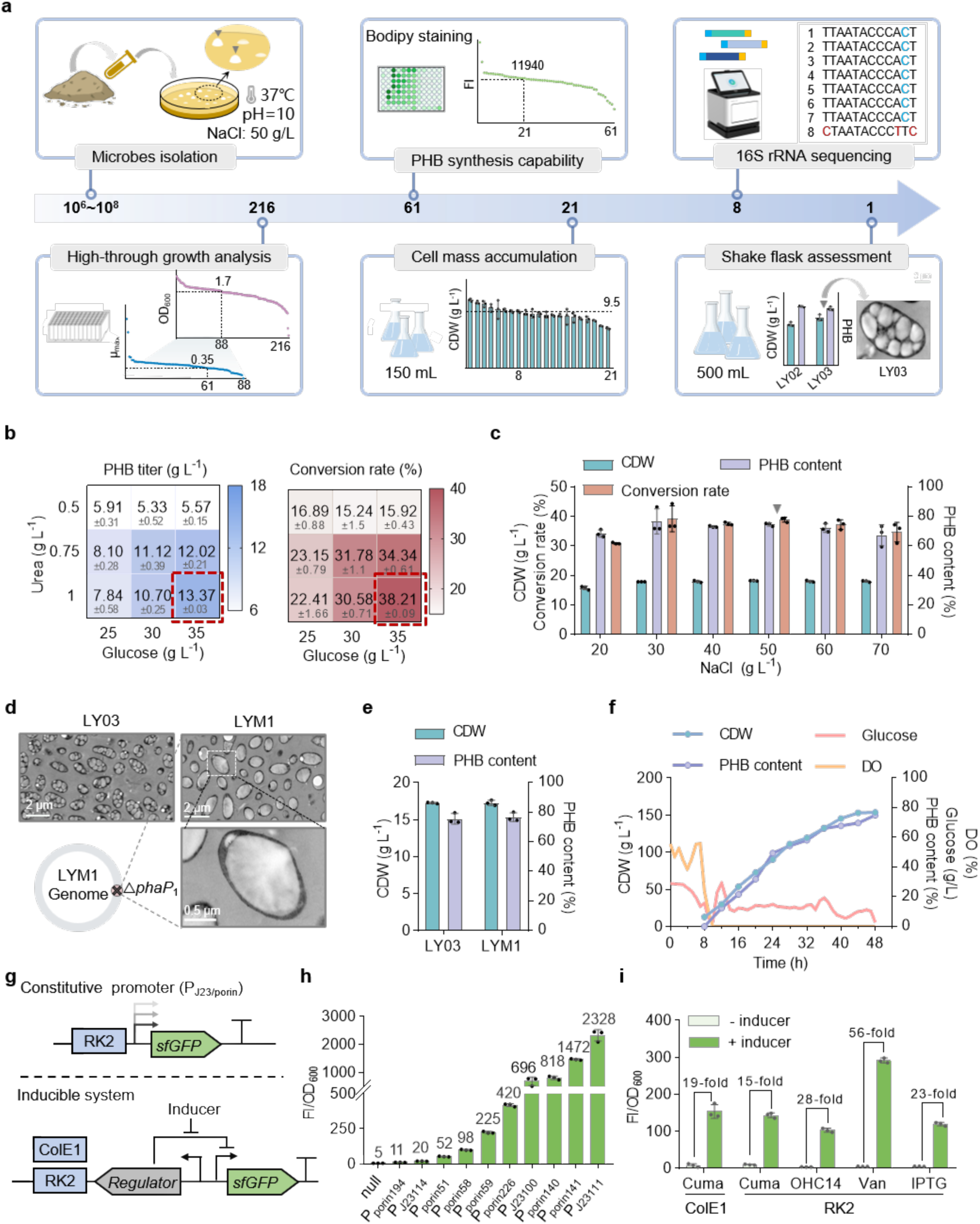
Isolation of high-performing halophilic chassis for PHA production. **(a)** Schematic workflow for isolating high-PHB-producing halophilic strain, *Halomonas* LY03 (LY03), from saline and alkaline environments in Xinjiang, China. Numbers within the arrows indicate the quantity of halophilic isolates from each screening step, including colony collection, high-through growth analysis, PHB synthesis assessment, cell growth evaluation in shake flask, 16S rRNA identification and shake flask study for PHB and cell mass (CDW) accumulation. Detailed screening criteria are provided in Fig. S1. **(b)** Shake flask study of carbon-to-nitrogen (C/N) ratio in 50MM medium that affects the PHB accumulation and glucose-to-PHB conversion rate by LY03. Dashed box in red indicates optimal C/N ratio (35 g/L glucose & 1 g/L urea, 50MMG). **(c)** Studying the effect of NaCl supplementation in 50MMG on the cell growth and PHB synthesis by LY03. **(d)** Transmission electron micrograph (TEM) analysis of intracellularly accumulated PHB granules by LY03 compared to its derivate with defected *phaP*_1_ gene related to PHB granule size control, namely LYM1 (LY03Δ*phaP*_1_). **(e)** Comparison of cell growth (CDW) and PHB content by LYM1 and LY03 grown in 50MMG for 48 h in a 150-mL shake flask. **(f)** Time-course profiles of CDW, PHB content, dissolved oxygen (DO) and residual glucose by LYM1 grown in a 5-L bioreactor using glucose as a feedstock. **(g)** Schema of constitutive and inducible promoter characterization in LYM1 using sfGFP as a reporter constructed on pSEVA341 (colE1 replicon) or pSEVA321 (RK2 replicon), respectively. **(h)** Normalized fluorescence intensity (FI/OD_600_) of sfGFP driven by various P_J23_ and P*_porin_* promoters. **(i)** Expression foldchange of sfGFP controlled by different inducible systems under saturated induction state (1 mM cuminic acid, 0.01 mM OHC14, 1 mM vanillic acid, and 10 mg L^−1^ IPTG) against the non-induction state. PHB titer (or content) and glucose conversion rate of shake flask studies were assessed after 48-h fermentation. For shake flask study, error bars represent standard deviations, n = 3. For fed-batch study, n = 1.

Additionally, the wild type *Halomonas* LY03 is no exception, which accumulates numerous tiny PHA granules inside cells, thereby complicates downstream separation and purification procedures^16, 62^. To address this limitation, we knocked out the granule-size-associated phasin-encoding gene *phaP*_1_^16^ to form the recombinant strain LYM1, which produces PHB with slightly higher molecular weight relative to a *phaP*_1_-deficient TD strain TDH4^16^ (Table S4). This mutant exclusively synthesizes large single PHA granule while fully retaining its native PHB biosynthesis capacity (Figs. 1d-1e). More remarkably, high-cell-density fed-batch cultivation of LYM1 was carried out in a 5-L bioreactor using glucose as a sole carbon source, achieving over 154 g L⁻¹ CDW containing 75 wt% PHB (Fig. 1f). Subsequently, for further metabolic engineering purposes, we systematically evaluated the performance of a panel of genetic parts and tools functional in LY03, including antibiotic resistance markers (Fig. S2c), expression vectors (Fig. 1g), constitutive promoters (Fig. 1h, Table S5), and chemical-induced systems (Fig. 1i, Fig. S5).

Furthermore, we assessed the capacity of LYM1 or its derivatives to utilize three representative non-grain carbon feedstocks of low cost, including tropical cassava starch (CS)^63^, waste soybean oil (WSO) from soy sauce manufacturing (Haitian Co., Ltd.), and crude glycerol (CG), a major by-product of biodiesel^64^, under both batch and fed-batch fermentation regimes. For CS utilization, a two-step enzymatic hydrolysis process was performed to obtain two hydrolysates, including directly-harvested crude hydrolysate (CHL) and its refined one after post-hydrolysis alkali treatment and centrifugation (HL). Expectedly, shake-flask study confirmed that LYM1 achieved comparable CDW and PHB content when grown on either CHL or HL compared to glucose. Followed by fed-batch test, a CDW of 83 g L⁻¹ containing 76 wt% PHB was obtained from HL only (Figs. 2a-2c). By contrast, LYM1 can directly utilize the saline WSO, mainly containing long chain-length fatty acids (Table S6), to yield up to 13 g L^−1^ PHB (17.6 g L⁻¹ CDW) after 48-h shake-flask cultivation under lower salt concentration (30 g L^−1^ NaCl), with an oil-to-PHB conversion rate reaching 43.3% (Figs. 2d-2f and Figs. S6c-6d). To avoid saponification under high pH, fed-batch study of CDW and PHB content by LYM1 was conducted in a 5-L bioreactor by maintain the pH at 7, achieving 54.8 g L⁻¹ CDW with high PHB content, reaching 80.5 wt% (Fig. 2g). Similarly, LYM1 manifests comparable CDW and PHB accumulation capacity whenever using pure (shake flask, 15.6 g L⁻¹ CDW & 69.5 wt% PHB; 5-L bioreactor, 73 g L⁻¹ CDW & 75.8 wt% PHB) or crude (shake flask, 14.4 g L⁻¹ CDW & 65.0 wt% PHB) glycerol (Fig. 2i, Figs. S7a-7b). Building on these promising results, we proceeded to engineer LYM1 for copolymer production consisting of 3HB and 3-hydroxypropionate (3HP), namely PHBP, from CG only, by redirecting the glycerol metabolism toward 3HP synthesis. Specifically, we introduced a heterologous glycerol-to-3-hydroxypropionaldehyde (3HPA) biosynthetic pathway mediated by the *dhaB-gdrAB* operon, which cooperated with endogenous *iolA*, a putative homolog of *pduP* encoding aldehyde dehydrogenase, to transform 3HPA into 3HP-CoA (Fig. 2h, Figs. S7d & S8, Table S7). Meanwhile, the competing 3HP degradation pathway governed by the *dddAC* gene cluster was knocked out (Fig. S7c). Via such combinatorial metabolic modifications, the engineered LYM1 strain successfully produced PHBP with a 3HP fraction of ∼8 mol% (Fig. 2j, Figs. S7e-7f). Scale-up test of PHBP was next validated in 5-L fed-batch fermentation using CG as a sole carbon source, achieving 36 g L⁻¹ CDW with 75.8 wt% PHBP (15 mol% 3HP) accumulation (Fig. 2k).

**Figure 2.**
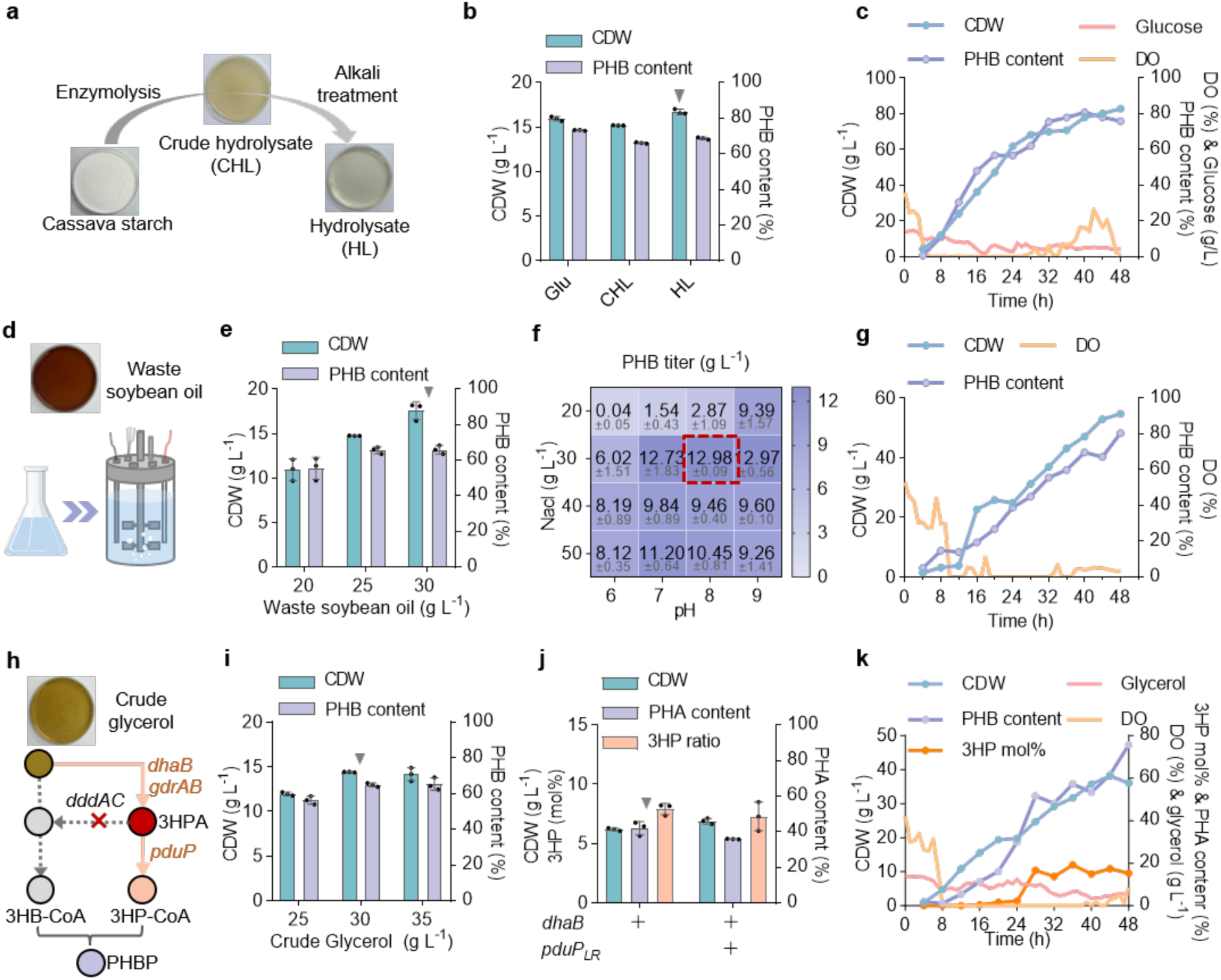
Developing LYM1 as a non-grain substrate-utilizing chassis for PHA production. **(a)** Hydrolysis workflow of cassava starch: a 600 g L^−1^ starch suspension is orderly digested by α-amylase (80 °C, 1.5 h) and glucoamylase (60 °C, 2 h) at pH=5 to obtain crude hydrolysate (CHL), followed by pH adjusted to 11 using 5 M NaOH and centrifugation to generate a purified hydrolysate (HL) containing approximately 540 g L^−1^ glucose after centrifugation (3580 ×g). **(b)** Shake flask study of CDW and PHB content by LYM1 grown on glucose (35 g L^−1^), CHL and HL (diluted to 35 g L^−1^ glucose), respectively, as a sole carbon source. **(c)** Fed-batch study of CDW, PHB content, DO and residual glucose by LYM1 conducted in a 5-L bioreactor using HL as feedstock only. **(d)** PHA production by LYM1 from waste soybean oil. **(e)** Shake flask study of CDW and PHB content by LYM1 grown in 50MM supplemented with 20, 25 and 30 g L^−1^ waste soybean oil. **(f)** Studying the effect of NaCl concentration and pH within the MM media on PHB accumulation by LYM1 grown on 30 g L^−1^ waste soybean oil as a sole carbon source. Dashed box in red indicates the optimal pH and NaCl condition for waste soybean oil-to-PHB bioconversion. **(g)** Fed-batch study of CDW, PHB content, DO and residual glucose by LYM1 conducted in a 5-L bioreactor using waste soybean oil as feedstock only. **(h)** Metabolic engineering of LYM1 for PHBP [P(3HB-*co*-3HP)] synthesis from glycerol. Arrows in light orange indicate heterologous 3HP-CoA synthesis pathway comprising *dhaB-gdrAB* cluster (encoding glycerol dehydratase and its activators from *Klebsiella pneumoniae* HS11286) and *pduP* gene (encoding CoA-propanoylating propanal dehydrogenase from *Lactobacillus reuteri* CF48-3A). The bypass of 3HPA encoded by *dddAC* operon was deleted to generate reinforced flux towards 3HP monomer synthesis. **(i)** Shake flask study of CDW and PHB content by LYM1 grown in 50MM supplemented with 25, 30 and 35 g L^−1^ crude glycerol, respectively, as a sole carbon source. **(j)** Studying the effect of heterologous expression of *dhaB-gdrAB* and *pduP*_ST_ module on PHBP synthesis (CDW, PHA content and 3HP molar ratio) in recombinant LYM1 grown on 30 g L^−1^ crude glycerol only. **(k)** Fed-batch study of CDW, PHA content, 3HP molar ratio, DO and residual glycerol by recombinant LYM1 harboring *dbaB-gdrAB* expression module conducted in a 5-L bioreactor using crude glycerol as feedstock only. For shake flask study, error bars represent standard deviations, n = 3. For fed-batch study, n = 1.

In summary, our findings established LY03 as a robust and versatile chassis for PHA biosynthesis, exemplifying promising potential of valorizing low-cost feedstocks for sustainable bioplastics via tailored genetic manipulation.

### Design of efficient genomic integration with high-level expression

As gene expression on chromosomal loci is generally significantly lower than that on mostly plasmid-based systems^65^, establishing efficient genomic integration strategies that enable high-level heterologous expression is critical for developing stable, antibiotic-free production strains^66^. Although several genome-editing approaches have been documented in *Halomonas* spp.^43, 44^, challenges such as from-plasmid-to-genome reduction on expression levels and low integration efficiency remain largely unsolved. Here, two strategies were established to address these limitations, including I) integrase/recombinase-mediated rapid genomic integration of large-fragment (> 9 kb) (Fig. 3a), and II) stop codon (TAA)-dependent enhancement of chromosomal gene expression (Fig. 3e).

**Figure 3.**
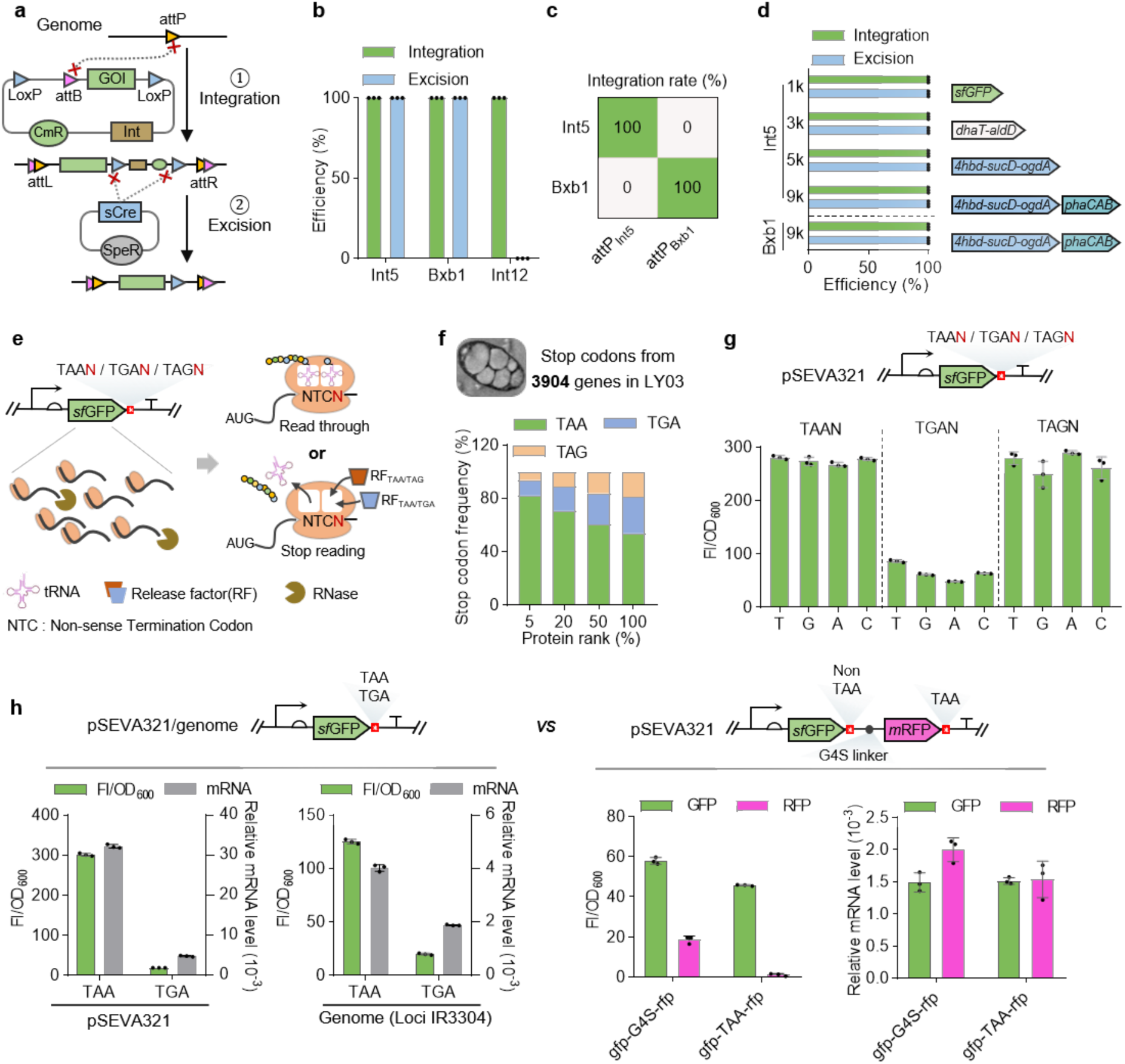
Developing recombinase-based genomic integration tools and stop codon-preference rationale for chromosomal gene expression. **(a)** Schema of two-step genomic integration mediated by integrase and recombinase. **(b)** Integration and excision efficiency of targeted integration module (P_J23110_-*sfgfp*) employing different integrase systems. Eight clones were randomly selected from each group for quantative efficiency evaluation. **(c)** Orthogonality assessment between Int5 and Bxb1 integrase systems. The ‘attP’ site of Int5 and Bxb1 was pre-integrated on loci IR_0815_ and IR_3304_, respectively, for further integration assay. **(d)** Integration and excision efficiency of different expression modules with increased fragment size across orthogonal integrase systems. **(e)** Schematic illustration of how stop codons function to terminate translation. **(f)** Distribution frequency of three stop codons (TAA, TAG and TGA) in different LYM1 gene sets grouped by protein expression levels (top 5%, 20%, 50% and 100%). Overall, 2914 out of 3904 genes (74.6%) were detected at the proteome level. **(g)** Normalized sfGFP fluorescence (FI/OD_600_) in LYM1 with different quadruplet stop codons (TAAN, TGAN, TAGN). (**h**) Normalized fluorescence (FI/OD_600_) and relative mRNA level of plasmid (pSEVA321)- and chromosome (IR_3304_)-based sfGFP terminated by TAA and TGA, respectively. Error bars represent standard deviations, n = 3.

For genomic integration, a donor plasmid was first constructed harboring the target cassette flanked by loxP sites, as well as a non-target region mainly consisting of an integrase gene (e.g., int5, int12, bxb1) and its recognition site ‘attB’. This plasmid was then integrated into a pre-defined genomic locus that had previously been modified to carry the attP site cognate with attB sequence using the homologous recombination method^43^. Following loxP-specific excision mediated by the sCre recombinase^51^, the non-target region was removed from the chromosome, leaving a clean insertion of the target cassette^48^. First, the integration efficiencies of three selected integrases (Int5, Int12, and Bxb1) were individually assessed by inserting a constitutive sfGFP expression cassette into each corresponding attP site pre-loaded in LYM1. All three integrases exhibited 100% integration efficiency as verified by colony PCR (Fig. 3b). However, in contrast to the complete sCre-mediated excision achieved in the Int5 and Bxb1 systems, no successful excision was detected for Int12 (Fig. 3b). Accordingly, the Int5- and Bxb1-based integration systems were further evaluated for orthogonality in an engineered LYM1 strain harboring attP_int5_ and attP_bxb1_ sites at the IR_0815_ and IR_3304_ loci, respectively (Fig. 3c). Expectedly, no cross talk was observed between Int5 and Bxb1, highlighting strong potential for simultaneous, dual-locus and site-specific genomic integration, particularly suitable for high-throughput tuning of dual-module pathways at the chromosomal level. Second, we constructed a series of expression cassettes spanning 1 kb to over 9 kb for evaluating the integration capacity of the Int5-based system. These modules, widely employed in *Halomonas* strains, included *sfgfp* (1.4 kb), *aldD-dhaT* (3.5 kb)^67^, *4hbd-sucD-ogdA* (5.1 kb)^68^, and *4hbd-sucD-ogdA-phaCAB* (9.1 kb). Among 8 randomly selected colonies, both integration and excision exhibited a 100% success rate (Fig. 3d). Furthermore, consistent with the Int5 system, the Bxb1 integrase was further validated to maintain high integration efficiency even for the largest cassette (9.1 kb) (Fig. 3d). Nevertheless, these results establish an integrase/recombinase-mediated genomic integration tool that enables rapid, highly efficient insertion of target cassettes across a range of sizes.

For single-gene expression design, numerous previous studies have focused on engineering the parts within the open reading frame (ORF)^69^, which typically consists sequentially of an insulating spacer^70^, promoter^71^, RiboJ^72^, RBS^73^, coding sequence (CDS), and terminator^74^. Similar studies have also been reported in *Halomonas* chassis^34^. Based on these knowledges, we herein further explore the potential impact of stop codons on gene expression levels in both LYM1 through genome-wide stop codon profiling (Fig. 3e). Genome-wide proteomic analysis of LYM1 grown on nitrogen-rich (5 g L⁻¹ urea) and nitrogen-poor (1 g L⁻¹ urea) conditions revealed that TAA serves as the stop codon for over 50% of the 3904 genes (Fig. 3f). Notably, this proportion rises sharply to 82% in genes with protein abundance ranked in the top 5% (Fig. 3f, Fig. S9, and Tables S8-S9). A consistent stop codon usage bias was further validated at the transcriptomic and translational levels (Fig. S9 and Tables S8-S9). These findings indicate that TAA-mediated translational termination is significantly enriched in highly expressed genes of LYM1, pointing to a potential evolutionary selection mechanism governing gene expression efficiency. Driven by this distinct TAA enrichment pattern, we experimentally investigated the effects of stop codon identity on protein expression levels. The expression of reporter gene *sfgfp* harboring three stop codons (TAA, TAG and TGA), along with their immediate single-base downstream contexts, was individually characterized in both LYM1 and *E. coli*. Consistent with our hypothesis, constructs carrying TAA exhibited substantially superior robustness in maintaining high-level protein expression compared with TAG and TGA counterparts (Fig. 3g, Fig. S10). We further quantified the mRNA and protein abundances of *sfgfp* with TAA and TGA stop codons on both plasmid (pSEVA321)- and chromosome (IR_3304_)-based systems in recombinant LYM1. The results confirmed that the TAA stop codon confers significantly elevated mRNA transcription and protein translation levels in contrast to TGA (Fig. 3h, left panel).

Generally, the variation of translational termination efficiency modulates gene expression via two primary mechanisms: inefficient termination can trigger ribosome stalling and consequently accelerate mRNA degradation, while unterminated readthrough sequesters working ribosomes and thereby diminishes downstream translational efficiency (Fig. 3e)^75, 76^. The elevated mRNA abundance observed for TAA-containing reporter constructs support the involvement of the former mechanism. To further investigate whether the TAA stop codon affects the full-length transcription efficiency when locating between two genes within a bi-cistron cluster, we constructed two dual-reporter systems, namely sfGFP-G4S-mRFP encoding fusion reporters and sfGFP-TAA-G4S-mRFP with TAA-based termination of sfGFP. Fluorescence quantification showed comparable sfGFP signals from two constructs, whereas mRFP expression was remarkably reduced upon TAA in sfGFP-TAA-G4S-mRFP group. However, the mRNA levels of both sfGFP and mRFP remained almost the same, implying robust translational termination of TAA with negligible transcription influence and readthrough activity (Fig. 3h, right panel). Therefore, these findings confirmed that the stop codon used for desired gene expression can significantly influence the translational readthrough activity and transcriptional level, thereby modulate the expression levels of target gene.

These results establish an efficient genomic integration tool and a stop codon (TAA)-dependent optimization strategy for enhanced gene expression in LYM1.

### Algorithm-guided mining of high-performing neutral integration site (HENIS)

Despite the high efficiency of integrase/recombinase-mediated genomic integration and TAA-guided enhancement of gene expression, chromosomally integrated genes still exhibited substantial expression variability and overall lower expression levels compared to plasmid-based systems. This limitation is probably attributed to intrinsic genomic position effects^52, 53, 77^ (Fig. 4a). To tackle this bottleneck, we developed an algorithm-guided toolbox termed SiteSeek, which enables the genome-wide identification of high-expression neutral integration sites (HENIS). SiteSeek integrates whole-genome annotations and transcriptomic profiles, alongside open-source databases including RegulonDB (https://regulondb.ccg.unam.mx)^78^, BDGP (https://www.fruitfly.org)^79^, etc., to screen and filter qualified intergenic regions (IGR). This pipeline excludes loci containing functional or regulatory elements, such as rRNA, tRNA, sRNA, tandem repeat framework (TRF), promoter, transcription factor binding site (TFBS, namely operator), etc. (Fig. 4b, Figs. S11-S13). Particularly, we developed a fast-blast k-mer alignment method for rapid identification of homologous operator sequences retrieved from the RegulonDB database, enabling efficient TFBS filtration within the upstream-yielded IGRs (Fig. S12). Combined with the above filtering criteria, this pipeline ultimately retains genomic segments neighbored by highly expressed genes, yielding high-quality HENIS candidates for robust gene integration (Figs. 4b-4c).

**Figure 4.**
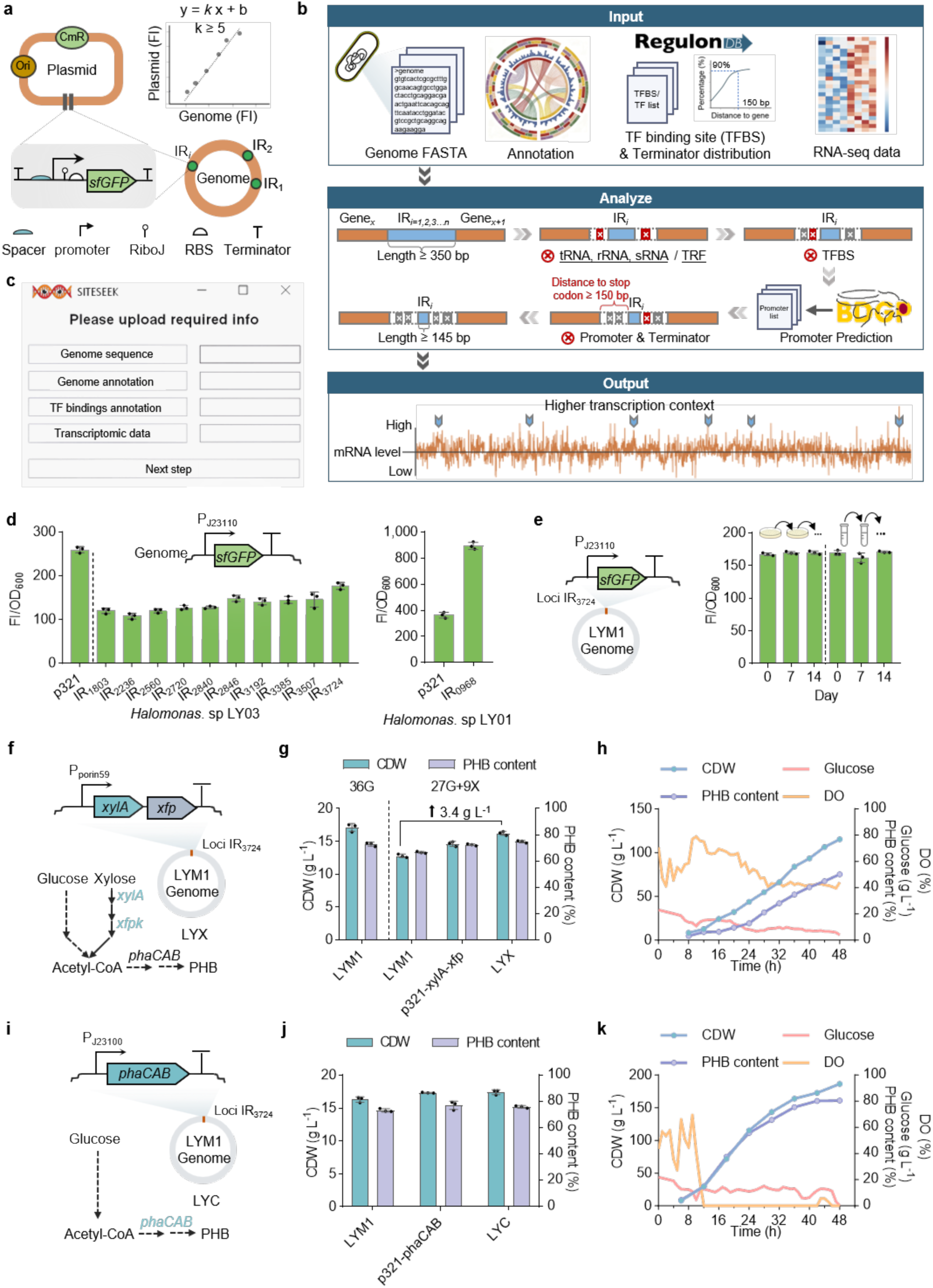
Algorithm-guided mining of high-expression neutral integration sites in LYM1. **(a)** Schematic illustration of gene expression variance across different expression systems, including various genomic loci on chromosome-carried system against plasmid-carried system. **(b)** Workflow for genome-wide high-expression neutral integration site (HENIS) screening. Parameters and required input data can be independently adjusted according to customized requirements and strain characteristics. **(c)** An user interface of ‘SiteSeek’, a toolkit for HENIS screening across different hosts. **(d)** sfGFP expression levels on different identified HENIS loci (see Table S11) in LYM1 and LY01, respectively, grown in 60LB medium compared to the control by relevant recombinant strains harboring sfGFP expression module (P_J23110_-*sfgfp*) constructed on pSEVA321. **(e)** Robustness assessment of integrated module (P_J23110_-*sfgfp*) on loci IR_3724_ in LYM1 via 14-day subculturing on a 60LB agar plate and in a 15-mL tube containing 5-mL 60LB medium. Normalized fluorescence recorded on day-0, -7 and -14 was determined by microplate reader. **(f)** Schematic design of chromosomally engineered LYM1 for PHB synthesis from glucose and xylose, namely LYX, which contains a fine-tuned xylose-utilizing module (P_porin59_-*xylA*-*xfp*) inserted on locus IR_3724_ in LYM1. **(g)** Shake flask study of LYX grown in 50MM supplemented with 27 g L^−1^ glucose and 9 g L^−1^ xylose compared to LYM1 (36 g L^−1^ glucose, or 27 g L^−1^ glucose & 9 g L^−1^ xylose) and its recombinant harboring expression vessel (p321-*xylA*-*xfp*) (27 g L^−1^ glucose & 9 g L^−1^ xylose). **(h)** Time-course profiles of CDW, PHB content, dissolved oxygen (DO) and residual glucose by LYX grown in a 5-L bioreactor using glucose and xylose (3:1, w/w) as feedstocks. **(i)** Schematic design of chromosomally engineered LYM1 for enhanced PHB synthesis, namely LYC, which contains a PHB synthesis module (P_J23100_-*phaCAB*) inserted on locus IR_3724_ in LYM1. **(j)** Shake flask study of LYC grown in 50MMG compared to LYM1 and its recombinant harboring expression vessel p321-*phaCAB*. **(k)** Time-course profiles of CDW, PHB content, dissolved oxygen (DO) and residual glucose by LYC grown in a 5-L bioreactor using glucose only. Error bars represent standard deviations, n = 3. For fed-batch studies in **h** and **k**, n = 1.

We then applied the SiteSeek pipeline to the LYM1 chassis and identified 10 HENIS candidates, all of which enabled successful sfGFP reporter integration using the integrase/recombinase-mediated genomic integration method (Tables S11-12). Fluorescence quantification showed that these ten chromosomal loci yielded normalized fluorescent levels ranging from 41% to 65% of the plasmid (pSEVA321)-carried system (Fig. 4d). This performance is substantially superior to previously characterized chromosomal sites in *Halomonas* TD01, which typically retain only ∼10% of the plasmid-based expression level^34^. To further validate the cross-strain portability and robustness of SiteSeek pipeline, we applied it to another *Halomonas* isolate, LY01 (GDMCC No: 62635), which is phylogenetically distant from LYM1 and TD01 and exhibits a native phenotype of single-granule PHA accumulation (Fig. S14, Table S11). A single high-confidence HENIS (IR_0968_) was identified and successfully integrated with the sfGFP reporter. Notably, this locus drove reporter expression level up to 246% over the pSEVA321 plasmid-based system (Fig. 4d), indicating potent intrinsic transcriptional activity and validating the reliable predictive capacity of SiteSeek across distinct genetic backgrounds. Additionally, long-term expression stability was assessed by recombinant LYM1 harboring the sfGFP expression module integrated at IR_3724_, the locus showing the highest expression level (∼65% of pSEVA321 plasmid) (Fig. 4d). Following 14-generation passaging growth on non-selective condition, colony PCR at the 7^th^ and 14^th^ generations verified stable retention of the integrated sfGFP cassette, and the normalized fluorescence intensity remained unchanged throughout passaging (Fig. 4e, Fig. S15). These results demonstrate that the IR_3724_ locus confers outstanding genetic robustness and high-level heterologous expression capacity at chromosomal level.

For functional gene expression uses of industrial purposes, two fine-tuned clusters encoding xylose utilization (*xylA-xfp* driven by P_porin59_) (Fig. 4f, Figs. S16a-16b)^80^ and PHB synthesis (phaCAB driven by P_J23100_) (Fig 4i) were integrated on IR_3724_ locus to yield the antibiotic-free LYX and LYC recombinants, respectively. For shake flask studies, the LYX grown on glucose-xylose mixture (27 g L⁻¹ glucose + 9 g L⁻¹ xylose) (Fig. S16c) and LYC strain grown on 35 g L⁻¹ glucose displayed 3.4 g L⁻¹ and 1.1 g L⁻¹ increase of CDW, respectively, compared to the start host LYM1 grown on corresponding conditions (Figs. 4g & 4j). Notably, higher CDW and PHB content can be observed by LYX in contrast to the recombinant LYM1 harboring *xylA-xfp* module on plasmid pSEVA321 (Fig. 4g). More importantly, scale-up fed-batch study of LYX and LYC conducted in a 5-L bioreactor yielded up to 115 g L⁻¹ (50 wt% PHB) and 186 g L⁻¹ (81 wt% PHB) CDW, respectively, which were significantly higher than that by LYM1 grown in the same conditions (45 g L⁻¹ CDW with 2.4 wt% PHB by LYX, 154 g L⁻¹ CDW with 75 wt% PHB by LYC) (Figs. 1f, 4h & 4k, Fig. S16d). These results demonstrate proven robustness of IR_3724_ locus serving as an effective HENIS for the chromosomal expression of desired genetic modules and exemplify successful cases in efficient non-grain sugar utilization and high-titer PHB biosynthesis.

### High-titer P34HB production by chromosomally engineered LYM1

In spite of the high-yield of PHB, the brittle nature is still the main bottleneck for widely commercial applications. Therefore, the introduction of non-side-chain monomers can tune the material property from hard to flexible and with enhanced processability suitable for wide-range applications. Herein, a biosynthesis platform for PHA copolymers consisting of 3HB and various α, ω-hydroxy acyl (α,ω-HA) units was constructed based on LYM1 by introducing a synthetic pathway encoding alcohol/aldehyde dehydrogenase (*dhaT_Pp_-aldD_Hb_* driven by P_porin58_, a high-strength promoter) and CoA transferase (*orfZ* driven by P_porin194_, a low-strength promoter) (Fig. 5a)^81^. Shake flask studies showed that the P34HB copolymer synthesized from glucose and C4 1,4-butanediol (1,4-BDO) as mixed carbon source exhibited highest levels of CDW (13.9 g L⁻¹) and PHA content (65 wt% containing 8.6 mol% 4HB) compared to other groups using C3, C5 and C6 diols (Fig. 5b, Fig. S17). The incorporation of various α, ω-HA units within different copolymers was respectively verified via ^1^H NMR assay (Fig. S17). Before genomic integration, different alcohol (2 homologies) and aldehyde (5 homologies) dehydrogenases from various hosts were constructed to form 10 combinations^82, 83, 84, 85^, of which the optimal recombinant (*ydcW_Kp_-dhaT_Pp_*) yielded the best overall performance, thereby used for genomic integration (Figs. 5c-5d). Therefore, a LYB recombinant was obtained by integrating *ydcW-dhaT* (requiring higher expression) and *orfZ* (requiring relatively lower expression) cluster at IR_3724_ and IR_1803_ loci, respectively, showing similar cell growth performance compared to PHB-producing host LYM1 with CDW, P34HB content and 4HB ratio reaching over 16.5 g L⁻¹, 74 wt% and 7.6 mol% (Figs. 5e-5f). Scale-up fed-batch fermentation of P34HB production by LYB was further validated in a 5-L bioreactor using glucose and 1,4-BDO as co-substrates, achieving up to 178 g L⁻¹ CDW containing 78 wt% P34HB content with a 4HB ratio of 5.5 mol% (Fig. 5g). Notably, the P34HB titer and productivity reached 139 g L^−1^ and 2.90 g L⁻¹ h⁻¹, respectively.

**Figure 5.**
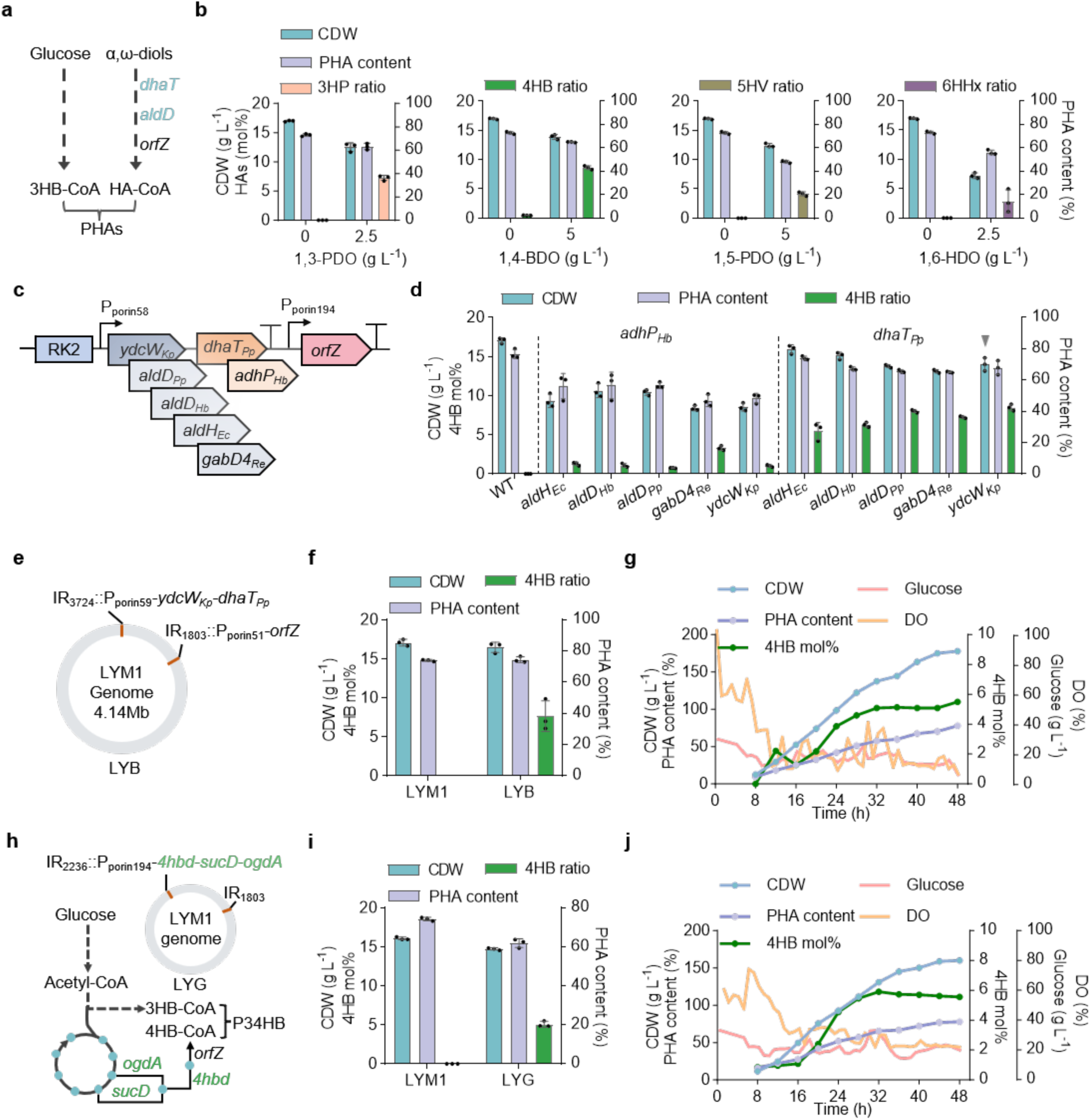
Engineering LYM1 for high-level P34HB production. **(a)** Schema for diol-derived PHA synthesis by introducing heterologous *dhaT*_Pp_, *aldD*_Pp_ and *orfZ* genes encoding alcohol dehydrogenase (*Pseudomonas putida* KT2440), aldehyde dehydrogenase (*Pseudomonas putida* KT2440), and CoA transferase (*Clostridium kluyveri* DSM 555), respectively. HA-CoA indicates different precursors, including 3-Hydroxypropionyl-CoA (3HP-CoA), 4-Hydroxybutyryl-CoA (4HB-CoA), 5-Hydroxypentanoyl-CoA (5HV-CoA), and 6-Hydroxyhexanoyl-CoA (6HHx-CoA), for different diol-derived PHA synthesis. **(b)** Shake flask study of different diol-derived PHA copolymers by recombinant LYM1 harboring *aldD-dhaT* and *orfZ* expression modules grown in 50MMG supplemented relevant structure-related carbon sources including 1,3-propanediol (1,3-PDO for 3HP), 1,4-butanediol (1,4-BDO for 4HB), 1,5-pentanediol (1,5-PDO for 5HV) or 1,6-hexanediol (1,6-HDO for 6HHx), respectively. **(c)** Pathway optimization for P34HB synthesis by screening different homologs of aldehyde dehydrogenase (5 candidates) and alcohol dehydrogenase (2 candidates). **(d)** Shake flask study of P34HB production by recombinant LYM1 harboring different combinations of *dhaT* and *aldD* homologies grown in 50MMG supplemented with 5 g L^−1^ 1,4-BDO. Grey inverted triangle indicates optimal combination. **(e)** Schema of constructing LYB strain by integrating P_porin59_-*ydcW*_Kp_*-dhaT*_Pp_ and P_porin51_-*orfZ* modules on loci IR_3724_ and IR_1803_, respectively. **(f)** Shake flask study of P34HB by LYB grown in 50MMG supplemented with 5 g L^−1^ 1,4-BDO. **(g)** Fed-batch study of P34HB production by LYB fed with glucose and 1,4-BDO conducted in a 5-L bioreactor. **(h)** Construction of LYG strain by integrating the de novo 4HB synthesis module encoded by *4hbd-sucD-ogdA* cluster (loci IR_2236_) and *orfZ* (loci IR_1803_). **(i)** Shake flask study of P34HB by LYG grown in 50MMG. **(j)** Fed-batch study of P34HB production by LYG conducted in a 5-L bioreactor using glucose as a sole carbon source. For shake flask studies, error bars represent standard deviations, n = 3. For fed-batch studies in **g** and **j**, n = 1.

To further eliminate the reliance on expensive precursor 1,4-BDO, we then integrated the fine-tuned *de novo* 4HB biosynthesis pathway composed of *4hbd-sucD-ogdA* and *orfZ* clusters^68^ on IR_2236_ and IR_1803,_ respectively, in LYM1, forming LYG strain (Fig. 5h). Similarly, the shake-flask fermentation showed robust cell growth by LYG in contrast to LYM1 with negligible decrease of CDW and 4HB molar ratio except the slight decrease of P34HB content reducing to 61.8 wt% (Fig. 5i). In 5-L fermentation, LYG achieved a CDW of 159 g L⁻¹ containing 77 wt% P34HB with a 4HB molar ratio of 5.6 mol% (Fig. 5j), maintaining comparable performance on P34HB titer and productivity compared to LYB strain.

In summary, the chromosomally engineered strains achieved robust, high-titer and antibiotic-free P34HB production in both batch and fed-batch conditions, demonstrating the high efficiency and precision of our genomic integration and expression tools for tailoring rapid and rational industrial strain development.

### Industrial-scale production of PHB and P34HB via fed-batch and two-stage continuous fermentation

Scaling up the biomanufacturing process from in-lab success (< 30 L), to pilot (200 L to 5 m^3^)- and industrial (> 5 m^3^)-scale validation with sufficient techno-economic feasibility is extremely critical to overcome the valley-of-death barrier that hinders the commercialization of most bioproducts. We first performed pilot-scale fermentation of two representative engineered strains, LYC and LYG, which enable high-yield production of PHB and P34HB, respectively, using glucose as the sole carbon source (Fig. 6a). In 2-m³ bioreactor cultivation, LYC attained a CDW of 163 g L⁻¹ with a PHB content of 82%, while LYG achieved a CDW of 149 g L⁻¹, accumulating 83 wt% P34HB with a 4HB molar ratio of 7.9 mol%. Consistent performance was further validated in three independent replicate fermentations in a 20-m³ industrial plant. At this larger scale, LYC reached 155 g L⁻¹ CDW with 78.2 wt% PHB, and LYG produced 162 g L⁻¹ CDW containing 78 wt% P34HB with a 6.1 mol% 4HB ratio. These results demonstrate the exceptional robustness and high-titer PHA accumulation capability of the engineered strains across pilot and industrial fermentation scales.

**Figure 6.**
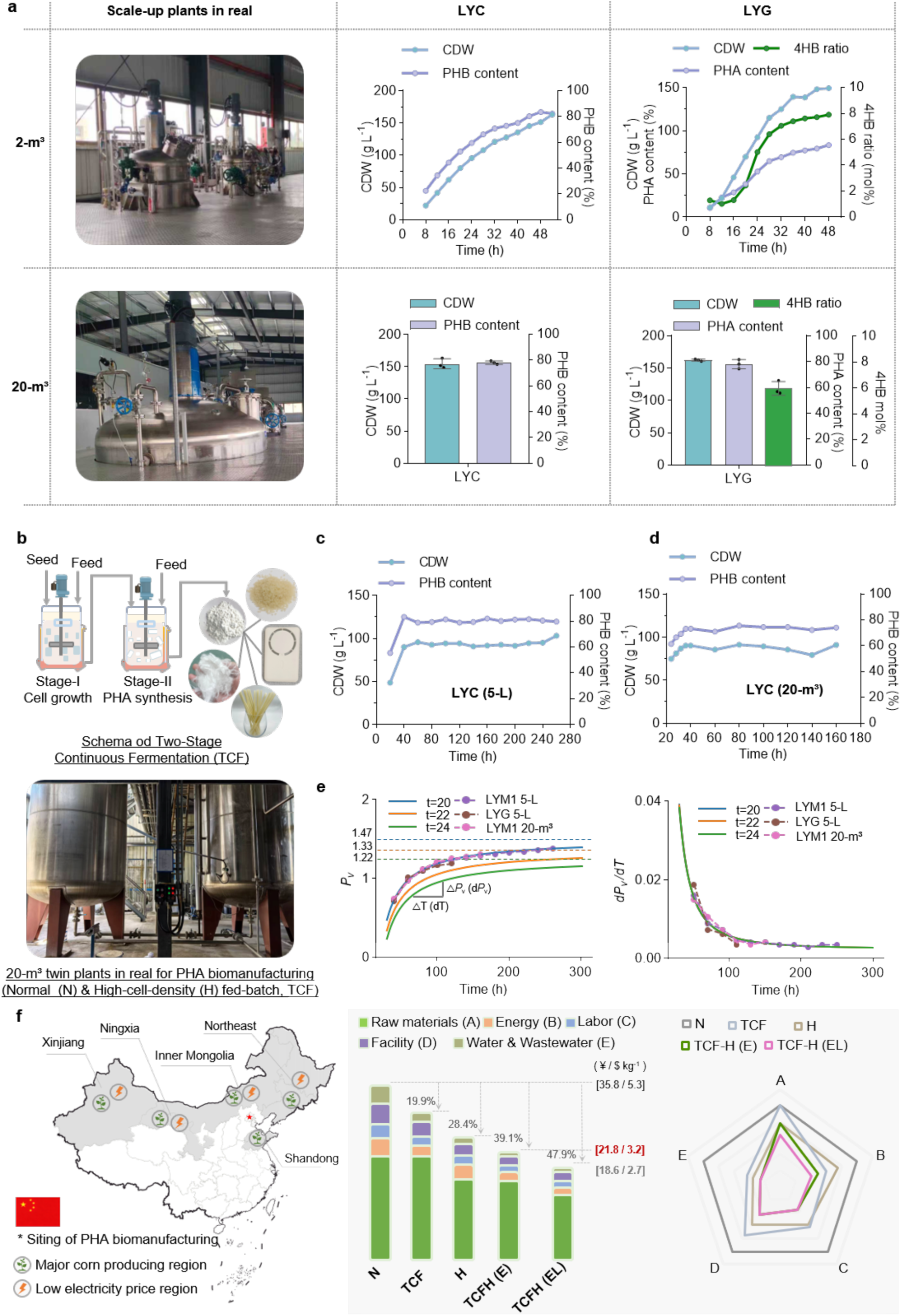
Fed-batch and two-stage continuous fermentation for scale-up PHA production by engineered LYM1. (**a**) High cell-density fed-batch fermentation of LYC and LYG conducted in a 2 m³ and 20 m³ bioreactor, respectively. **(b)** Schematic of a two-stage continuous fermentation (TCF) process for PHA production. Stage I mainly supports the cell growth with true cell mass accumulation for high-cell-density seed generation under nitrogen-rich condition (20 h per cycle), whereas Stage II switches into PHA synthesis under nitrogen-limited condition, followed by periodic harvest of fermentation broth for downstream extraction and purification for end-product manufacturing. A representative industrial-scale (20-m³) bioreactor is shown below. **(c-d)** Two-stage continuous fermentation for PHB production by LYC conducted in a 5-L **(c)** and 20-m³ **(d)** bioreactor. Cell cultures were harvested every 20 h for CDW and PHB content analysis. **(e)** Production-efficiency-per-volume ratio (*P*_v_) of two-stage continuous fermentation versus fed-batch fermentation as a function of total biomanufacturing time (T) (left panel) and its first derivative (d*P*_v_/dT) (right panel). Solid curves show computed values at harvest intervals (t) of 20, 22 and 24 h, whereas dashed curves show experimental data points fitted to the computation ones. The curves gradually flatten and approach the theoretical optimal limits of both *P*_v_ and d*P*_v_/dT when T tends to infinity. Detailed calculations are described in Supplementary Note. **(f)** Techno-economic analysis (TEA) of different fermentation configurations for PHA production. The geographic distribution of low-cost maize-derived glucose and electricity in China identifies large-scale favorable locations for plant deployment. Stacked cost breakdown and radar analysis of normal fed-batch (N), two-stage continuous (TCF), high-cell-density fed-batch (H), high-cell-density TCF under estimated (TCFH-E) and location-optimized TCFH-E (TCFH-EL) scenarios. All costs were generated from the TEA model (see Supplementary Data 1). For fed-batch fermentation, n = 1 (2-m³) and n = 3 (20-m³); for two-stage continuous fermentation, n = 1.

Fed-batch fermentation has long been the dominant manufacturing process for intracellular PHA production, which inherently restricts further improvements in productivity and cost-effectiveness during batch-to-batch operation. To address this bottleneck, we developed a two-stage continuous fermentation (TCF) strategy consisting of a high-nitrogen cell growth stage and a nitrogen-limited PHA biosynthesis stage (Fig. 6b). This design enables the cyclic and orderly accumulation of biomass and PHA with minimal manual operation, greatly improving the continuity and efficiency of industrial PHA fermentation^57^. We first validated the feasibility of this design in a 5-L twin bioreactor system using LYC and LYG strains conducted under an unoptimized normal cell-density fermentation process, which were continuously cultured for 260 and 120 h, respectively (Fig. 6c, Fig. S18). For LYC, 12 consecutive harvests (every 20 h) yielded stable CDW of 90-100 g L⁻¹ with a consistent PHB content of approximately 80 wt% PHB accumulation, demonstrating robust long-term production stability (Fig. 6c). Similarly, LYG exhibited steady performance across 5 harvest cycles (every 20 h), maintaining a CDW of 77-95 g L⁻¹ containing 67-75 wt% P34HB with a 4HB molar ratio of 5-7 mol% (Fig. S18). Encouraged by the success in 5-L scale, we further scaled up the TCF process of LYC to an industrial scale (20-m^3^) twin-plant system. After 160-h continuous operation, 7 successive harvests in total were achieved in every 20 h with CDW and PHB content reaching around 80-90 g L⁻¹ and ∼75 wt% (Fig. 6d), respectively, demonstrating promising operational stability, process scalability and industrial feasibility of the newly established TCF strategy for high-efficiency PHA biomanufacturing.

To quantitatively evaluate the time- and space-utilization efficiency of different fermentation modes, we defined a volumetric production efficiency ratio (*P_v_*) (Supplementary Note). This metric benchmarks the overall operational efficiency of the total bioreactor volume throughout the entire fermentation duration by comparing the TCF strategy with conventional normal fed-batch fermentation (N, ∼100 g L^−1^ CDW). Unlike TCF processes, fed-batch workflows involve inevitable batch-to-batch downtime intervals, which substantially increase capital investment and labor costs while restricting continuous production output. At the 5-L scale with comparable PHA synthesis performance, the TCF strategy improved the *P_v_* values by 35% and 16% for LYC and LYG strains, respectively (Fig. 6e, left panel). A 27% *P_v_* enhancement was also achieved by LYC in 20-m³ pilot-scale TCF fermentation (Fig. 6e, left panel). The shortened harvest cycles inherent to the TCF system account for the elevated *P_v_* performance, reflecting improved equipment utilization and reduced operational and fixed-asset costs. Furthermore, the first derivative of *P_v_* with respect to fermentation time (d*P_v_/*d*T*) revealed that the marginal economic benefit of TCF gradually reaches to a theoretical optimal level over extended operation, becoming economically viable when the fermentation duration exceeds 140 h or even longer (Fig. 6e, right panel).

To systematically assess the techno-economic feasibility of fed-batch and TCF fermentation modes, we established a comprehensive techno-economic assessment (TEA) model that integrates multi-dimensional cost components, including raw materials, energy consumption, labor, facility depreciation, water usage and wastewater treatment (Fig. 6f, Supplementary Data 1). The normal fed-batch fermentation exhibited a PHA production cost of approximately ¥35.8 kg⁻¹ ($5.3 kg⁻¹), which aligns well with previously reported values for microbial PHA manufacturing^61^. In contrast, the TCF process reduced the production cost to ¥28.7 kg⁻¹ ($4.2 kg⁻¹) under the same fermentation performance, primarily attributed to enhanced volumetric utilization of fermentation facilities. Further economic optimization was projected by integrating high-cell-density cultivation (≥ 150 g L^−1^ CDW containing over 80 wt% PHA that independently contributes 28.4% cost reduction compared to the ‘N’ group; Figs. 4h, 4k, 5g, 5j, 6a & 6f) into the TCF framework (TCFH (E), E denotes estimated scenario), which could lower the production cost to ¥21.8 kg⁻¹ ($3.2 kg⁻¹). This value is considerably lower than that of standalone high-cell-density fed-batch fermentation (¥25.6 kg⁻¹, or $3.8 kg⁻¹), benefited from the synergistically improved *P_v_* and reduced raw material consumption. In addition, a location-dependent industrial deployment strategy was proposed to leverage the low-cost resource endowments in regions including Xinjiang, Gansu, Inner Mongolia, Shandong province and Northeast China. This optimized scenario (TCFH (EL), EL denotes estimated low-cost region) further cuts production costs by 15%, reaching ¥18.6 kg⁻¹ ($2.7 kg⁻¹) and approaching the commercial price of polylactic acid (PLA).

## Discussion

Despite decades of intensive research and industrial endeavors to accelerate PHA commercialization, the persistent drawbacks of high production cost and suboptimal material properties, especially for PHB, still impede the market competitiveness of PHA when compared to the traditional petrol-based plastics and mainstream degradable polyesters like PLA, PBAT, PBS, etc. As a type of rising-star chassis for PHA synthesis, the *Halomonas* strains^8^ exhibit several superior characteristics unattainable by conventional chassis strains, including extremely low contamination risk^33^, robust scalability^41, 86^, reduced energy consumption^17^, and simplified process-control complexity^87^. However, considerable room remains for further optimization, especially for sufficient tool enabling precise plasmid-to-chromosome integration^88^, achieving higher-level PHA titer and productivity^89^, before *Halomonas* can truly become a versatile all-round performer for industrial-scale PHAs. In this study, we tailored a begin-with-an-end-in-mind engineering pipeline to facilitate the isolate-to-industry translation for cost-effective PHA production based on the high-performing halophilic isolate, *Halomonas* LY03.

Compared to the previously reported *Halomonas* chassis strains, LY03 supports higher cell-density growth, stronger alkaline tolerance, broader feedstock utilization and superior growth robustness against drastic fluctuations in cultivation conditions (Figs. 1a-1f, 2, 6a, Figs. S1, 3f, 4). These physiological traits significantly advance the production level of PHAs (e.g., PHB, P34HB) with the highest titer exceeding 150 g L^−1^ under non-sterile, open fed-batch condition. More importantly, multidimensional tools for precise genomic integrative expression developed herein can significantly boost the strain development efficiency by directly transferring a plasmid-based fine-tuned expression module onto a predictable HENIS locus whatever in *Halomonas* hosts like LY03 and LY01, or other microbial chassis including *E. coli*, *Pseudomonas* (data not showed), etc. This circumvents the need for high-throughput screening, which typically relies on expensive equipment and involves uncertain, lengthy R&D cycles. Additionally, the successful breakthrough of two-stage continuous PHA fermentation at both lab (5-L) and industry (20 m^3^) scales implies a more stable and efficient PHA-manufacturing paradigm allowing improved productivity and material property stability in sharp contrast to the conventional fed-batch process (Figs. 6b-6d, Table S13). Further process optimization by integrating high cell-density fermentation into the TCF process system is expected to bring substantial improvements in both production efficiency and capacity, representing one of the most promising strategies to achieve cost reductions far beyond current industrial levels (Fig. 6f, Supplementary Note).

As a newly isolated chassis, further efforts are still urgently required to improve the utilization efficiency of non-grain (or waste-derived) feedstocks to industrially relevant levels, and to expand the PHA product spectrum with enhanced material properties by incorporating diverse monomers beyond 3HB and α,ω-HA units like 3HP, 4HB, 5HV, and 6HHx (Fig. 5, Fig. S17). Notably, extended exploration was implemented to achieve high-titer P34HB (∼113 g L^−1^) production with over 10 mol% 4HB, conferring significantly improved flexibility than the ones containing 5-6 mol% only, at 20-m^3^ scale fermentation by fine-tuning the fermentation process and chromosomal expression level of 4HB-synthesis pathway (Fig. S19). Besides, a preliminary investigation was performed to convert waste soybean oil (WSO) into the medium-chain-length copolymer PHBHHx, composed of 3HB and 3-hydroxyhexanoate (3HHx), by introducing heterologous *phaC* and *phaJ* genes in the *phaP*_1_-deficient strain LYM1. Prototype production of PHBHHx with a 3HHx ratio of 3-to-4 mol% was thus achieved (Fig. S6), highlighting the promising potential and feasibility of engineering the LY03 chassis for the synthesis of diverse functional PHA materials with tailored monomer compositions in future studies^90, 91, 92, 93^.

In summary, our study successfully isolated and engineered a halophilic chassis for scalable high-titer PHA productions with strong cost competitiveness. Meanwhile, the development of precise genomic integration tools and a two-stage continuous fermentation process lays solid foundations for PHA industrialization, and offers valuable rationales for chromosomal expression design applicable to different microbial chassis, and a novel-type fermentation paradigm for long-term continuously biomanufacturing of intracellular products.

## Methods and Materials

### Plasmids, strains, and culture media

General molecular biology techniques like PCR amplification, gel extraction, multi-fragment DNA ligation (e.g., Gibson assembly) and sequencing were used for plasmid construction. DNA primers synthesis and Sanger sequencing were performed by Sangon Biotech (China). High-fidelity DNA and Taq polymerase were purchased from Genestar (China) and Vazyme (China), respectively. Gel extraction and purification were performed using a DNA Gel Extraction Kit from Tiangen Biotech (China). DNA fragments were assembled using the EZ-HiFi Seamless Cloning Kit (Genestar, China). All procedures were carried out according to the manufacturers’ protocols unless otherwise specified.

All bacterial strains and plasmids used in this study were listed in Tables S1 and S2. *Escherichia coli* S17-1, the plasmid-constructing host and donor cell for conjugation, was cultivated in LB medium (10 g L⁻¹ NaCl, 10 g L⁻¹ tryptone, and 5 g L⁻¹ yeast extract) at 37 °C under 220 rpm. *Halomonas* hosts and their recombinants were cultured at 37 °C under 220 rpm in 60LB medium (derived from LB containing 60 g L⁻¹ NaCl) or 50MM medium.

The generally used 50MM medium was composed of 50 g L⁻¹ NaCl, 1 g L⁻¹ yeast extract, 35 g L⁻¹ glucose, 1 g L⁻¹ urea, 0.2 g L⁻¹ MgSO₄, 5 g L⁻¹ Na₂HPO₄·12H₂O, 0.75 g L⁻¹ KH₂PO₄, 10 mL L⁻¹ trace element solution I, and 1 mL L⁻¹ trace element solution II. Trace element solution I contained 2 g L⁻¹ CaCl₂ and 5 g L⁻¹ Fe(III)-NH₄-citrate dissolved in 1 M HCl. Trace element solution II contained 300 mg L⁻¹ H₃BO₃, 200 mg L⁻¹ CoCl₂·12H₂O, 10 mg L⁻¹ CuSO₄·5H₂O, 100 mg L⁻¹ ZnSO₄·7H₂O, 30 mg L⁻¹ MnCl₂·4H₂O, 20 mg L⁻¹ NiCl₂·6H₂O, and 30 mg L⁻¹ Na₂MoO₄·2H₂O. The pH of the culture medium for *Halomonas* cells was adjusted to 9.0–10.0 using 5 M NaOH before inoculation.

### Halophilic chassis screening for high-yield PHA production

Firstly, saline and alkaline mud samples were collected from Xinjiang province and suspended in lab using a 60LB medium. After filtering by the sterile gauze, the obtained filtrate was diluted to 10^2^-to10^5^-fold and spread on a 50MMG agar plate (pH = 10) for 48-h incubation at 37 °C. Colonies with larger size (diameter > 1mm), rounder-shape morphology, and poor transparency (especially for the white and opaque ones) were selected as candidates for high PHA-producing isolates.

Secondly, single colony of each isolate was inoculated into 1 mL 60LB medium in a 96-deep-well plate for 12-h growth at 37 °C (1000 rpm). The pre-cultured seed cells were 1 vol% inoculated into a fresh 60LB medium for 12-h growth under the same condition. Each culture was then 100-fold diluted into 200 μL 50MMG medium in a 96-well microplate for online monitoring of optical density at 600 nm (OD₆₀₀) every 30 mins over 24-h growth at 37 °C (600 rpm) in a multi-function microplate reader (Thermo Fisher Scientific, USA). The growth rate of each selected isolate was calculated based on the obtained growth curve: μ = (OD₆₀₀(t_x+1_) – OD₆₀₀(t_x_)) / OD₆₀₀(t_x_), t_x_ indicates the time point at any time throughout the growth process. To assess the PHA-producing performance, cell cultures were obtained following the same procedure except the step that isolates grew 1-mL 50MMG medium was conducted in a 96-deep-well plate for 48-h culturing at 37 °C using a deep-well plate oscillator (Allsheng, China). The obtained cell cultures were diluted 20-fold (OD₆₀₀ ≈ 0.6–1.0), chilled on ice for 10 mins, and then centrifuged at 3000 × g for 5 min at 4 °C. Cell pellets were harvested and washed once with TSE buffer (10 mM Tris-HCl, 0.25 M sucrose, 1 mM EDTA, pH 7.5), incubated on ice for 10 min, and centrifuged again. The harvested cell pellets were resuspended with distilled water (1 mL) and stained with 5 μg/mL BODIPY 493/503 (Cayman Chemical) in the dark for 5 min. Finally, the strained cells were washed with distilled water again for single-cell fluorescent intensity (FI) measurement at excitation/emission wavelengths of 488/510 nm using a multi-function microplate reader (Thermo Fisher Scientific, USA). The isolates with normalized FI (FI/OD_600_) over 11940 were selected for shake-flask fermentation in a 150-mL conical flask (48 h, 220 rpm) for biomass accumulation (cell dry weight, CDW) evaluation. Subsequently, isolates with CDW over 9.5 g L^−1^ were selected for 16S rRNA sequencing (Sangon Biotech, Shanghai, China), which was amplified using the general primers (forward: 5′-AGAGTTTGATCCTGGCTCAG-3′); reverse: (5′-ACGGCTACCTTGTTACGACT-3′). The obtained 16S rRNA sequence of each isolate was aligned to wipe off reductant duplications for phylogenetic analysis using MEGA software (Version 11). Finally, the isolates with distinct 16S rRNA were delivered for shake flask fermentation (48 h, 220 rpm) in a 500-mL conical flask containing 50-mL 50MMG medium under at 37 °C.

### Shake-flask fermentation

The 1^st^ seed culture was prepared using a single colony as an inoculum grown in 5-mL 60LB medium (37 °C, 220 rpm) for 12 h. Followed by 1 vol% inoculation of 1^st^ seed culture into 20-mL fresh 60LB medium for 12-h growth, a 2^nd^ seed culture was obtained and used as a final inoculum (5 vol%) to grow in 20- or 50-mL 50MMG medium. Antibiotics (50 μg mL^−1^ kanamycin, 25 μg mL^−1^ chloramphenicol, 100 μg mL^−1^ spectinomycin, and 100 μg mL^−1^ ampicillin) and required inducers (e.g., cumic acid, vanillic acid, IPTG, or 3-OHC14-HSL) were added into the medium whenever needed. After 48-h cultivation at 37 °C (220 rpm), cells were harvested and washed for CDW and PHA content analysis.

### Fed-batch fermentation

Seed cultures were prepared using the same procedure as described for shake-flask fermentation except the 2^nd^ seed culture was 5-to-10 vol% inoculated into the bioreactors (5 L, 2 m^3^, or 20 m^3^). All fermentations were initiated at a working volume of 35-to-65 vol% total capacity using a modified 50MMG medium containing 2 g L^−1^ urea, 12 g L^−1^ yeast extract (or waste corn steep liquor with similar organic nutrient), and 20 g L^−1^ glucose. The temperature and pH were maintained at 37°C and 8.5, respectively, during the fermentation process. Meanwhile, the agitation speed (rpm) and aeration (vvm, volume per volume per min) were adjusted to maintain the dissolved oxygen over 30% until their upper limits were reached. Three- to four-phase feedstocks were one-by-one fed to maintain the residual glucose (or glycerol) approximately at 5-10 g L^−1^ unless otherwise specified. For fed-batch fermentation using waste soybean oil as carbon source, the pH was adjusted at 7.0 and the NaCl concentration was decreased to 30 g L^−1^. Other operating parameters remained consistent across all fermentation studies unless otherwise noted.

### Two-stage continuous fermentation

Two-stage continuous fermentation process was designed and carried out in twin bioreactors (5-L and 20-m^3^). Specifically, the first-stage fermentation was initiated under nitrogen-rich condition to facilitate true cell mass accumulation with negligible PHA accumulation, thereby generating high-density cell culture (approximately 20 h), which would be transferred into another bioreactor for 20-h second-stage fermentation aiming to accumulate PHA under nitrogen-limited condition. About 5-to-10 vol% of first-stage cell culture was kept in the bioreactor used as an inoculum for following fermentation cycle after supplementing the same volume of basal medium as the initial condition. Residual glucose was maintained at 5-10 g L^−1^ throughout the two-stage fermentation process. The two-stage fermentation process last at least 260 h and 120 h long for PHB and P34HB production, respectively. Detailed calculations of the volumetric production efficiency ratio (*Pv*) are described in the Supplementary Note.

### Preparation of cassava starch hydrolysate

Cassava starch hydrolysate (CSH) was prepared as follows (per liter): 300 g of cassava starch was sampled and suspended in 500-mL distilled water and adjusted to pH at 5.0 using 5 M HCl. First, 12 mL of high-temperature α-amylase (Annzyme, AHA-200T) was added to the suspension and hydrolyzed at 80 °C for 30 min. An additional 300 g of cassava starch and 12 mL of α-amylase were then supplemented, the total hydrolysis volume was adjusted to 1 L, and hydrolysis was continued for another 1 h. Followed by cooling to 60 °C, 60 mL of glucoamylase (Annzyme, GA-260) was added into the hydrolysate for 2-h saccharification. Finally, the obtained crude hydrolysate was centrifuged at 8601 × g for 5 min to collect the supernatant as crude hydrolysate, namely CSL. The pellet was harvested and added into the next-round hydrolysis. Notably, the starch-releasing proteins in CSL can be optionally removed by adjusting the pH to 11 with 5 M NaOH, followed by incubation at room temperature for 2 h. The final purified CSL can be obtained after 5-min centrifugation at 8601 × g.

### Fluorescence characterization

Single colony was inoculated into 5-mL 60LB medium and pre-cultured at 37 °C (220 rpm) for 12 h. The pre-culture was subsequently 1% (v/v) inoculated into 1-mL 60LB in a 96-deep-well plate for 12-h growth at 37 °C (1000 rpm) in a microplate incubator (Allsheng, China). Then, the cell culture was passaged once following identical procedure for the next 24-h cultivation. Inducers were initially added into the medium whenever needed. Subsequently, cell culture was diluted 10-fold using phosphate-buffered saline (PBS), and 200 μL dilution was then transferred into a well in a 96-well plate for fluorescence measurement using a multi-function plate reader (Thermo Fisher Scientific, USA) with excitation and emission wavelengths set to 488 and 510 nm, respectively. Fluorescence intensity was normalized to optical density at 600 nm (OD_600_), namely FI/OD_600_.

### Conjugation

Targeted plasmids were transformed into *Halomonas* strains via conjugation using *E. coli* S17-1 as a donor. Donor and recipient strains were cultured separately for 8 to 12 h in LB and 60LB medium, respectively. Equal volumes of both cultures were harvested by centrifugation at 1500 × g for 2 min at 4 °C, and washed once using fresh 20LB medium (10 g L^−1^ tryptone, 5 g L^−1^ yeast extract, 20 g L^−1^ NaCl). The obtained cell pellets were individually resuspended and 1:1 (vol/vol) mixed together to generate cell resuspension, which was spread (60–80 μL) on an antibiotic-free 20LB agar plate for 6 to 12 h incubation at 37 °C. The resulting bacterial lawn was resuspended using fresh 60LB medium and spread (100–200 μL) on a 60LB agar plate containing appropriate antibiotics (generally Cm + Amp). Single colony of transconjugant can be obtained after 36 to 48 h incubation at 37 °C, followed by colony PCR verification.

### Genome editing

Gene deletion and recombinase-targeted ‘attP’ sequence insertion were achieved using a previously reported homologous recombination strategy based on the instable vector pRE112^43^ using ∼1 kb homology arms flanking the target locus. Site-specific integration of targeted genetic modules was then achieved through serine recombinase-mediated recombination, which was reported in previous studies in different hosts^94^.

By employing these approaches, *phaP*_1_ gene was deleted from the parental strain LY03 to generate recombinant LYM1. And attP_int5_ and attP_bxb1_ were respectively integrated into the genome of LYM1 at different loci via homologous recombination to obtain two recombinants, LYM3 (IR_1803_ for attP_int5_ and IR_3724_ for attP_bxb1_) and LYM4 (at IR_1803_ for attP_int5_ and IR_2236_ for attP_bxb1_). Based on these chassis, various chromosomally engineered recombinants were further constructed: LYX, *xylA-xfp* under the control of P_porin59_ on IR_3724_; LYC, *phaCAB* driven by P_J23100_ on IR_3724_; LYB, *ydcW_Kp_-dhaT_Pp_* driven by P_porin59_ on IR_3724_ and *orfZ* driven by P_porin51_ on IR_1803_; LYG, *4hbd-sucD-ogdA* driven by P_porin194_ on IR_2236_ and *orfZ* driven by P_porin51_ on IR_1803_.

### Genome Sequencing and Omics Profiling

Cell culture of LY03 or its derivates were collected via centrifugation at 3,000 × g for 10 min at 4 °C and washed twice with ice-cold ddH₂O, followed by storage at –80 °C. Genomic DNA was extracted using TIANamp Bacteria DNA toolkit (Tiangen, China) and sent to BGI Genomics (Shenzhen, China) for whole-genome sequencing. Whole-genome annotation was conducted using different database (e.g., RAST, KEGG, UNIPROT, etc.). For transcriptomic and proteomic analysis, frozen cell pellets were stored in liquid nitrogen and directly sent to Novogene Co., Ltd. (Beijing, China) for relative quantification (RPKM, reads per kilobase per million mapped reads) of detectable reads matched to annotated coding sequence.

### High-expression neutral integration site (HENIS) prediction by SiteSeek

HENIS regions were mined using a general software package, namely SiteSeek, depending on different input data, including genome sequence and annotation, regulon info, and transcriptome. Specifically, intergenic regions (IGRs) of at least 350 bp (adjustable parameter) (target_length) between two neighboring coding sequences (CDS) were first extracted. Secondly, IGRs containing annotated rRNA, tRNA, sRNA, or tandem repeat regions were removed to eliminate the potential effects of integrated module on non-coding functional elements. Thirdly, sequences of predicted transcription factor binding sites (TFBSs), promoters, and terminators within the IGR regions were deleted to avoid unexpected transcriptional interference. Specifically, TFBSs sequences were predicted using an optimized k-mer algorithm^95^ referring to the annotated regulon sequences of at least 8-bp with ≥ 90% sequence identity (adjustable parameter) compared to the ones generated from RegulonDB (*E. coli*). For each matched hit of TFBS, the aligned segment and its 10-bp flanking sequences were removed from the IGRs. Promoter regions were predicted using the neural network model in BDGP^79^ with a threshold of 0.9 (adjustable parameter). Sequences extending from each predicted promoter to the nearest downstream start codon (with an additional 10-bp buffer) were then trimmed. To define terminator regions, 317 experimentally validated terminators from RegulonDB were analyzed, showing that 90% are located within 150 bp downstream of the stop codon of the coding sequences (Fig. S13). Therefore, appropriate sequence deletion was further required when any IGR was less than 150 bp (adjustable parameter) away from the nearest TAA codon. After one-by-one filtering, a functional element-free IGRs library with sequence length over 145 bp (adjustable parameter) (min_segment) was collected. Finally, RNA-seq data were used to determine the transcription activities (Intensity, %) of both upstream and downstream flanking genes of each IGR candidate. The final HENIS library can be obtained by determining the adjustable parameters (e.g., target_length, min_segment, Intensity, etc. within the software interface). All parameter settings are provided in Tables S10-S11.

### Transmission electron microscopy (TEM) analysis

Cells were harvested by centrifugation at 1500 rpm for 3-5 min, washed twice with PBS, and fixed overnight at 4 °C using 2.5% glutaraldehyde prepared in PBS. After fixation, all subsequent steps, including post-fixation, dehydration, embedding, ultrathin sectioning, staining, and imaging, were performed by a commercial EM service provider (zkbaice, China). Samples were processed and visualized using a Tecnai G2 Spirit transmission electron microscope (FEI, USA) operated at 100 kV.

### Nuclear magnetic resonance (NMR) analysis

Extracted PHA materials (5–10 mg) were dissolved in deuterated chloroform (CDCl₃). And the ^1^H NMR spectra were recorded on an AVANCE III HD 400 MHz spectrometer (Bruker, Germany). Chemical shifts were reported in parts per million (ppm) relative to tetramethyl silane (TMS, 0.03% v/v) as the internal standard.

### Analytical methods and statistics

To determine cell dry weight (CDW, g L^−1^), 15 mL of each cell culture was centrifuged at 9000 × g for 5 min, washed once with distilled water to harvest cell pellets. After pre-freezing at −80 °C for 1 h, the obtained cell pellets were freeze-dried for 24 h (CREATRUST, China) for dry weight measurement, which can be used to calculate CDW of each sample.

For PHA quantification, ∼30 mg of each freeze-dried pellet was subjected to methanolysis in the mixture of 2-mL esterification solution (3% H₂SO₄ and 0.1% benzoic acid in methanol) and 2 mL chloroform at 99 °C for 4 h. After cooling to room temperature, 1-mL distilled water was added into the methanolysate, followed by 2-min vortexing (1500 rpm) and then phase-separation extraction for 2 h. The heavy organic phase was sampled (1 mL) and analyzed by gas chromatography (GC-2010 Pro, Shimadzu, Japan) using an SH-5 column (30 m × 0.25 mm ID, 0.1 μm film thickness). The standards of PHB, γ-butyrolactone, and 3HV (Sigma-Aldrich) were used for determining the content and molar ratio of 3HB, 4HB, and 3HV monomers, respectively.

The concentration of glucose and 3-hydroxypropionate (3HP) were measured using an LC-20 system (Shimadzu, Japan) equipped with an Aminex HPX-87H column (Bio-Rad, USA) and a RID-10A refractive index detector. Cell culture was firstly centrifuged at 12,000 × g for 5 min to obtain supernatant, which was then diluted 10-fold and filterted using a 0.22 μm membrane to prepare sample for HPLC analysis. An external standard curve with five concentration gradients was constructed first for quantification of each analyte.

## Data availability

All data supporting the findings of this study are available within the Article and its Supplementary Information. Additional materials, including transcription factor binding site annotations from RegulonDB, genome annotations and transcriptomic data of *Halomonas* LY03 and LY01, as well as source-code files for the SiteSeek, are also available whenever needed with the permission of MedPHA Bioscience Co., Ltd. Source data are provided with this paper.

## Acknowledgements

We are grateful to Prof. Jing Huang (East China Normal University/China) and Prof. Guo-Qiang Chen (Tsinghua University/China) for kindly donating the pSEVA series plasmids and *Halomonas* TD strain (used as a control in this study), respectively. This project was financially supported by grants from the National Natural Science Foundation of China (Grant No. 32322003 to YJW, No. 32501308 to LYN), the Fundamental Research Funds for the Central Universities (Grant No. 2025ZYGXZR096 to YJW), Department of Science and Technology of Guangdong Province (No. 2022B1111080006) and Guangdong Basic and Applied Basic Research Foundation (No. 2026A1515011148 to LYN).

## Author contributions

J.W.Y. conceived the study, and P.L., and X.X.Y. wrote the paper. P.L., X.X.Y., Y.H.D., C.W., H.Y., Y.X.L. performed the experiments. Z.F.L. implemented the computational process for SiteSeek. W.ST. and H.W.S. supervised the scale-up and continuous fermentation process. L.S.Y., J.Y.L., Y.C.X., and L.L.Z. participated in the research. J.W.Y. and Y.N.L. supervised the study. J.W.Y. revised the paper. All authors contributed to the article have approved the submission.

## References

1. Raza, Z. A., Abid, S. & Banat, I. M. Polyhydroxyalkanoates: characteristics, production, recent developments and applications. Int. Biodeterior. Biodegrad. 126, 45–56 (2018).

2. Sullivan, K. P., et al. Mixed plastics waste valorization through tandem chemical oxidation and biological funneling. Science 378, 207–211 (2022).

3. Mai, J., et al. Synthesis and physical properties of polyhydroxyalkanoate (PHA)-based block copolymers: A review. Int. J. Biol. Macromol. 263, 130204 (2024).

4. Chen, G. Q. & Patel, M. K. Plastics derived from biological sources: present and future: a technical and environmental review. Chem. Rev. 112, 2082–2099 (2012).

5. Choi, J. I., Lee, S. Y. & Han, K. Cloning of the *Alcaligenes latus* polyhydroxyalkanoate biosynthesis genes and use of these genes for enhanced production of Poly(3-hydroxybutyrate) in *Escherichia coli*. Appl. Environ. Microbiol. 64, 4897–4903 (1998).

6. Jiang, T., et al. Enhancing oil feedstock utilization for high-yield low-carbon polyhydroxyalkanoates industrial bioproduction. Metab. Eng. 91, 44–58 (2025).

7. Kahar, P., Tsuge, T., Taguchi, K. & Doi, Y. High yield production of polyhydroxyalkanoates from soybean oil by *Ralstonia eutropha* and its recombinant strain. Polym. Degrad. Stab. 83, 79–86 (2004).

8. Tan, D., Xue, Y. S., Aibaidula, G. & Chen, G. Q. Unsterile and continuous production of polyhydroxybutyrate by *Halomonas* TD01. Bioresour. Technol. 102, 8130–8136 (2011).

9. Ryu, H. W., Hahn, S. K., Chang, Y. K. & Chang, H. N. Production of poly(3-hydroxybutyrate) by high cell density fed-batch culture of *Alcaligenes eutrophus* with phospate limitation. Biotechnol. Bioeng. 55, 28–32 (1997).

10. Muigano, M. N., Mauti, G. O., Anami, S. E. & Onguso, J. M. Advances and challenges in polyhydroxyalkanoates (PHA) production using *Halomonas* species: A review. Int. J. Biol. Macromol. 309, 142850 (2025).

11. Aytar Celik, P., et al. A novel higher polyhydroxybutyrate producer *Halomonas halmophila* 18H with unique cell factory attributes. Bioresour. Technol. 372, 128669 (2023).

12. Hammami, K., et al. Extremophilic bacterium *Halomonas* desertis G11 as a cell factory for poly-3-hydroxybutyrate-*co-*3-hydroxyvalerate copolymer’s production. Front. Bioeng. Biotechnol.10, 878843 (2022).

13. Wang, M. R., Tao, S., Lin, N., Song, X. & Li, Z. J. Development of *Photobacterium* sp. LN01 as a versatile halophilic platform for tunable biosynthesis of Polyhydroxyalkanoates. ACS Synth. Biol.14, 4090–4099 (2025).

14. Chen, Y. L., et al. Cell cizes matter for industrial bioproduction, a case of polyhydroxybutyrate. Adv. Sci. 12, e2412256 (2025).

15. Jiang, X. R., Yao, Z. H. & Chen, G. Q. Controlling cell volume for efficient PHB production by *Halomonas*. Metab. Eng. 44, 30–37 (2017).

16. Shen, R., Ning, Z. Y., Lan, Y. X., Chen, J. C. & Chen, G. Q. Manipulation of polyhydroxyalkanoate granular sizes in *Halomonas bluephagenesis*. Metab. Eng. 54, 117–126 (2019).

17. Ling, C., et al. Engineering NADH/NAD(+) ratio in *Halomonas bluephagenesis* for enhanced production of polyhydroxyalkanoates (PHA). Metab. Eng. 49, 275–286 (2018).

18. Ouyang, P., et al. Increasing oxygen availability for improving poly(3-hydroxybutyrate) production by *Halomonas*. Metab. Eng. 45, 20–31 (2018).

19. Yi, X., et al. Protein phase separation for enhanced production of 3-hydroxypropionate and polyhydroxybutyrate by *Halomonas*. Metab. Eng. 96, 302–312 (2026).

20. Yin, J., Yang, J., Yu, X., Chen, T. & He, S. Enhanced poly(3-hydroxybutyrate-*co*-3-hydroxyvalerate) production from high-concentration propionate by a novel halophile *Halomonas* sp. YJ01: Detoxification of the 2-methylcitrate cycle. Bioresour. Technol. 388, 129738 (2023).

21. Ji, M., et al. PHB production from food waste hydrolysates by *Halomonas bluephagenesis* harboring PHB operon linked with an essential gene. Metab. Eng. 77, 12–20 (2023).

22. Pernicova, I., et al. Production of polyhydroxyalkanoates on waste frying oil employing selected *Halomonas* strains. Bioresour. Technol. 292, 122028 (2019).

23. Benesova, P., Kucera, D., Marova, I. & Obruca, S. Chicken feather hydrolysate as an inexpensive complex nitrogen source for PHA production by *Cupriavidus necator* on waste frying oils. Lett. Appl. Microbiol. 65, 182–188 (2017).

24. Yuan, Y., et al. Polyhydroxyalkanoate production by engineered *Halomonas* grown in lignocellulose hydrolysate. Bioresour. Technol. 425, 132313 (2025).

25. Wang, Y., Hu, Y. S., Hao, T. & Li, Y. Q. An alkali-halophilic laccase facilitates highly efficient conversion of lignin into polyhydroxybutyrate. Bioresour. Technol. 433, 132728 (2025).

26. Luo, C. B., et al. Efficiently unsterile polyhydroxyalkanoate production from lignocellulose by using alkali-halophilic *Halomonas alkalicola* M2. Bioresour. Technol. 351, 126919 (2022).

27. Kourilova, X., Novackova, I., Koller, M. & Obruca, S. Evaluation of mesophilic *Burkholderia sacchari*, thermophilic *Schlegelella thermodepolymerans* and halophilic *Halomonas halophila* for polyhydroxyalkanoates production on model media mimicking lignocellulose hydrolysates. Bioresour. Technol. 325, 124704 (2021).

28. Sohn, Y. J., Son, J., Lim, H. J., Lim, S. H. & Park, S. J. Valorization of lignocellulosic biomass for polyhydroxyalkanoate production: Status and perspectives. Bioresour. Technol. 360, 127575 (2022).

29. Shin, Y., et al. Production of polyhydroxybutyrate by halotolerant *Halomonas cerina* YK44 using sugarcane molasses and soybean flour in tap water. Int. J. Biol. Macromol. 279, 135358 (2024).

30. Kulkarni, S. O., Kanekar, P. P., Jog, J. P., Sarnaik, S. S. & Nilegaonkar, S. S. Production of copolymer, poly (hydroxybutyrate-*co*-hydroxyvalerate) by *Halomonas campisalis* MCM B-1027 using agro-wastes. Int. J. Biol. Macromol. 72, 784–789 (2015).

31. Kim, B., et al. Polyhydroxybutyrate production from crude glycerol using a highly robust bacterial strain *Halomonas* sp. YLGW01. Int. J. Biol. Macromol. 236, 123997 (2023).

32. Ben Abdallah, M., et al. Advances in polyhydroxyalkanoate (PHA) production from renewable waste materials using halophilic microorganisms: A comprehensive review. Sci. Total Environ.963, 178452 (2025).

33. Chen, X., et al. Engineering *Halomonas bluephagenesis* TD01 for non-sterile production of poly(3-hydroxybutyrate-*co*-4-hydroxybutyrate). Bioresour. Technol. 244, 534–541 (2017).

34. Ye, J., et al. Stimulus response-based fine-tuning of polyhydroxyalkanoate pathway in *Halomonas*. Metab. Eng. 57, 85–95 (2020).

35. Tsuji, A., Takei, Y. & Azuma, Y. Establishment of genetic tools for genomic DNA engineering of *Halomonas* sp. KM-1, a bacterium with potential for biochemical production. Microb. Cell Fact.21, 122 (2022).

36. Liu, C., Yue, Y., Xue, Y., Zhou, C. & Ma, Y. CRISPR-Cas9 assisted non-homologous end joining genome editing system of *Halomonas bluephagenesis* for large DNA fragment deletion. Microb. Cell Fact. 22, 211 (2023).

37. Zhang, Y. H., et al. Engineering complex phenotypes in *Halomonas bluephagenesis* TD01 via large-fragment manipulation and multiplex base editing. Metab. Eng. 96, 405–419 (2026).

38. Wang, M. R., Yi, J. & Li, Z. J. Tailored base editing toolkits for functional genomics and metabolic engineering in the halophile *Salinivibrio*. J. Biotechnol. 416, 56–63 (2026).

39. Zhang, X., Lin, Y., Wu, Q., Wang, Y. & Chen, G. Q. Synthetic biology and genome-editing tools for improving PHA metabolic engineering. Trends Biotechnol. 38, 689–700 (2020).

40. Choi, S. Y., et al. Metabolic engineering for the synthesis of polyesters: A 100-year journey from polyhydroxyalkanoates to non-natural microbial polyesters. Metab. Eng. 58, 47–81 (2020).

41. Wen, R., et al. Pilot production of P(3HB-*co*-4HB) by engineered *Halomonas bluephagenesis* harboring an endogenous plasmid grown on glucose. Metab. Eng. 94, 153–168 (2026).

42. Wang, M.-R., Song, X., Tao, S. & Li, Z.-J. Systematic metabolic engineering of *Photobacterium* sp. TLY01 for high-yield biosynthesis of poly(3-hydroxybutyrate-*co*-4-hydroxybutyrate). Adv. Ind. Eng. Polym. Res. 9, 115–124 (2026).

43. Fu, X. Z., et al. Development of *Halomonas* TD01 as a host for open production of chemicals. Metab. Eng. 23, 78–91 (2014).

44. Qin, Q., et al. CRISPR/Cas9 editing genome of extremophile *Halomonas* spp. Metab. Eng. 47, 219–229 (2018).

45. Yan, Q. & Fong, S. S. Challenges and advances for genetic engineering of non-model bacteria and uses in consolidated bioprocessing. Front. Microbiol. 8, 2060 (2017).

46. Corts, A., Thomason, L. C., Costantino, N. & Court, D. L. Recombineering in non-Model bacteria. Curr. Protoc. 2, e605 (2022).

47. Brown, W. R., Lee, N. C., Xu, Z. & Smith, M. C. Serine recombinases as tools for genome engineering. Methods 53, 372–379 (2011).

48. Fogg, P. C., Colloms, S., Rosser, S., Stark, M. & Smith, M. C. New applications for phage integrases. J. Mol. Biol. 426, 2703–2716 (2014).

49. Snoeck, N., et al. Serine integrase recombinational engineering (SIRE): a versatile toolbox for genome editing. Biotechnol. Bioeng. 116, 364–374 (2019).

50. Hwang, S., Joung, C., Kim, W., Palsson, B. & Cho, B.-K. Recent advances in non-model bacterial chassis construction. Curr. Opin. Syst. Biol. 36, 100471 (2023).

51. Elmore, J. R., et al. High-throughput genetic engineering of nonmodel and undomesticated bacteria via iterative site-specific genome integration. Sci. Adv. 9, eade1285 (2023).

52. Goormans, A. R., et al. Comprehensive study on *Escherichia coli* genomic expression: does position really matter? Metab. Eng. 62, 10–19 (2020).

53. Cai, P., et al. Recombination machinery engineering facilitates metabolic engineering of the industrial yeast *Pichia pastoris*. Nucleic Acids Res. 49, 7791–7805 (2021).

54. Yu, W., Gao, J., Zhai, X. & Zhou, Y. J. Screening neutral sites for metabolic engineering of methylotrophic yeast *Ogataea polymorpha*. Synth. Syst. Biotechnol. 6, 63–68 (2021).

55. Kong, S., Yu, W., Gao, N., Zhai, X. & Zhou, Y. J. Expanding the neutral sites for integrated gene expression in *Saccharomyces cerevisiae*. FEMS Microbiol. Lett. 369, (2022).

56. Gao, J., Gao, N., Zhai, X. & Zhou, Y. J. Recombination machinery engineering for precise genome editing in methylotrophic yeast *Ogataea polymorpha*. iScience 24, 102168 (2021).

57. Blunt, W., Levin, D. B. & Cicek, N. Bioreactor operating strategies for improved polyhydroxyalkanoate (PHA) productivity. Polymers 10, 1197 (2018).

58. Atlić, A., et al. Continuous production of poly([R]-3-hydroxybutyrate) by *Cupriavidus necator* in a multistage bioreactor cascade. Appl. Microbiol. Biotechnol. 91, 295–304 (2011).

59. Lee, J. H., et al. Quantified high-throughput screening of *Escherichia coli* producing poly (3-hydroxybutyrate) based on FACS. Appl. Biochem. Biotechnol. 170, 1767–1779 (2013).

60. Kacmar, J., Carlson, R., Balogh, S. J. & Srienc, F. Staining and quantification of poly-3-hydroxybutyrate in *Saccharomyces cerevisiae* and *Cupriavidus necator* cell populations using automated flow cytometry. Cytometry A 69, 27–35 (2006).

61. Zhang, L., et al. Engineering low-salt growth *Halomonas Bluephagenesis* for cost-effective bioproduction combined with adaptive evolution. Metab. Eng. 79, 146–158 (2023).

62. Anderson, A. J. & Dawes, E. A. Occurrence, metabolism, metabolic role, and industrial uses of bacterial polyhydroxyalkanoates. Microbiol. Rev. 54, 450–472 (1990).

63. Huang, J., et al. Production of n-butanol from cassava bagasse hydrolysate by engineered *Clostridium tyrobutyricum* overexpressing adhE2: kinetics and cost analysis. Bioresour. Technol.292, 121969 (2019).

64. Kumar, L. R., Yellapu, S. K., Tyagi, R. D. & Zhang, X. A review on variation in crude glycerol composition, bio-valorization of crude and purified glycerol as carbon source for lipid production. Bioresour. Technol. 293, 122155 (2019).

65. Wang, Z. K., et al. Improving the intensity of integrated expression for microbial production. ACS Synth. Biol. 10, 2796–2807 (2021).

66. Li, L., Liu, X., Wei, K., Lu, Y. & Jiang, W. Synthetic biology approaches for chromosomal integration of genes and pathways in industrial microbial systems. Biotechnol. Adv. 37, 730–745 (2019).

67. Zhang, L., et al. Effective production of Poly(3-hydroxybutyrate-*co*-4-hydroxybutyrate) by engineered *Halomonas bluephagenesis* grown on glucose and 1,4-Butanediol. Bioresour. Technol. 355, 127270 (2022).

68. Ye, J., et al. Engineering of *Halomonas bluephagenesis* for low cost production of poly(3-hydroxybutyrate-*co*-4-hydroxybutyrate) from glucose. Metab. Eng. 47, 143–152 (2018).

69. Smanski, M. J., et al. Synthetic biology to access and expand nature’s chemical diversity. Nat. Rev. Microbiol. 14, 135–149 (2016).

70. Zong, Y., et al. Insulated transcriptional elements enable precise design of genetic circuits. Nat. Commun. 8, 52 (2017).

71. Mutalik, V. K., et al. Precise and reliable gene expression via standard transcription and translation initiation elements. Nat. Methods 10, 354–360 (2013).

72. Lou, C., Stanton, B., Chen, Y. J., Munsky, B. & Voigt, C. A. Ribozyme-based insulator parts buffer synthetic circuits from genetic context. Nat. Biotechnol. 30, 1137–1142 (2012).

73. Salis, H. M., Mirsky, E. A. & Voigt, C. A. Automated design of synthetic ribosome binding sites to control protein expression. Nat. Biotechnol. 27, 946–950 (2009).

74. Chen, Y. J., et al. Characterization of 582 natural and synthetic terminators and quantification of their design constraints. Nat. Methods 10, 659–664 (2013).

75. Dulebohn, D., Choy, J., Sundermeier, T., Okan, N. & Karzai, A. W. Trans-translation: the tmRNA-mediated surveillance mechanism for ribosome rescue, directed protein degradation, and nonstop mRNA decay. Biochemistry 46, 4681–4693 (2007).

76. Baggett, N. E., Zhang, Y. & Gross, C. A. Global analysis of translation termination in *E. coli*. PLoS Genet. 13, e1006676 (2017).

77. Bryant, J. A., Sellars, L. E., Busby, S. J. & Lee, D. J. Chromosome position effects on gene expression in *Escherichia coli* K-12. Nucleic Acids Res. 42, 11383–11392 (2014).

78. Salgado, H., et al. RegulonDB v12.0: a comprehensive resource of transcriptional regulation in *E. coli* K-12. Nucleic Acids Res. 52, D255–D264 (2024).

79. Reese, M. G. Application of a time-delay neural network to promoter annotation in the *Drosophila melanogaster* genome. Comput. Chem. 26, 51–56 (2001).

80. Liu, Y., et al. Metabolic engineering of *Halomonas bluephagenesis* for the production of ethylene glycol and glycolate from xylose. J. Biotechnol. 396, 36–40 (2024).

81. Yan, X., et al. Biosynthesis of diverse α, ω-diol-derived polyhydroxyalkanoates by engineered *Halomonas bluephagenesis*. Metab. Eng. 72, 275–288 (2022).

82. Jo, J.-E., et al. Cloning, expression, and characterization of an aldehyde dehydrogenase from *Escherichia coli* K-12 that utilizes 3-hydroxypropionaldehyde as a substrate. Appl. Microbiol. Biotechnol. 81, 51–60 (2008).

83. Honjo, H., Tsuruno, K., Tatsuke, T., Sato, M. & Hanai, T. Dual synthetic pathway for 3-hydroxypropionic acid production in engineered *Escherichia coli*. J. Biosci. Bioeng. 120, 199–204 (2015).

84. Kim, J. W., Ko, Y. S., Chae, T. U. & Lee, S. Y. High-level production of 3-hydroxypropionic acid from glycerol as a sole carbon source using metabolically engineered *Escherichia coli*. Biotechnol. Bioeng. 117, 2139–2152 (2020).

85. Jiang, X. R., Yan, X., Yu, L. P., Liu, X. Y. & Chen, G. Q. Hyperproduction of 3-hydroxypropionate by *Halomonas bluephagenesis*. Nat. Commun. 12, 1513 (2021).

86. Ye, J., et al. Pilot scale-up of poly(3-hydroxybutyrate-*co*-4-hydroxybutyrate) production by *Halomonas bluephagenesis* via cell growth adapted optimization process. Biotechnol. J. 13, e1800074 (2018).

87. Wang, S., et al. Establishment of low-cost production platforms of polyhydroxyalkanoate bioplastics from *Halomonas cupida* J9. Biotechnol. Bioeng. 121, 2106–2120 (2024).

88. Coimbra, A. A. B., Prakash, S., Jiménez, J. I. & Rios-Solis, L. Establishing *Halomonas* as a chassis for industrial biotechnology: advances in synthetic biology tool development and metabolic engineering strategies. Microb. Cell Fact. 24, 133 (2025).

89. Tan, D., Wang, Y., Tong, Y. & Chen, G. Q. Grand challenges for industrializing polyhydroxyalkanoates (PHAs). Trends Biotechnol. 39, 953–963 (2021).

90. He, H., et al. Engineering *Halomonas bluephagenesis* for pilot production of terpolymers containing 3-hydroxybutyrate, 4-hydroxybutyrate and 3-hydroxyvalerate from glucose. Metab. Eng. 90, 117–128 (2025).

91. Wang, H., et al. Production and characterization of copolymers consisting of 3-hydroxybutyrate and increased 3-hydroxyvalerate by β-oxidation weakened *Halomonas*. Metab. Eng. 89, 97–107 (2025).

92. Min Song, H., et al. Production of polyhydroxyalkanoates containing monomers conferring amorphous and elastomeric properties from renewable resources: current status and future perspectives. Bioresour. Technol. 366, 128114 (2022).

93. Yu, H. E., Choi, S. Y., Song, S., Park, S. J. & Lee, S. Y. High-level production of short- and medium-chain-length polyhydroxyalkanoates from glucose using metabolically engineered *Escherichia coli*. ChemSusChem 19, e202502029 (2026).

94. Hirano, N., Muroi, T., Takahashi, H. & Haruki, M. Site-specific recombinases as tools for heterologous gene integration. Appl. Microbiol. Biotechnol. 92, 227–239 (2011).

95. Šošic, M. & Šikic, M. Edlib: a C/C ++ library for fast, exact sequence alignment using edit distance. Bioinformatics 33, 1394–1395 (2017).

