## Supplementary material for "Tailoring Precise Genomic Integration Toward Isolate-to-Industry Strain Development for Scalable High-Titer Production of Polyhydroxyalkanoate": Table S1-S3, Figure S1-S9 and Supplementary Note

### Supplementary Tables

### Table S1. Strains used in this study.

| **Strains** | **Descriptions** | **References** |
| --- | --- | --- |
| *E. coli* S17-1 | A vector donor used for conjugation harbors the *tra* genes from plasmid RP4 in the chromosome | ^1^ |
| *Halomonas* TD01 | Wild-type *Halomonas* TD01 strain isolated from Aydingkol Lake in Xinjiang Province, China | ^2^ |
| *Halomonas* TDH4 | TD01 derivate with *orfZ* gene integration (G4 loci) and *phaP*_1_ gene deletion | ^3^ |
| *Halomonas* LY01 | Wild-type *Halomonas* LY01 strain isolated from Turpan in Xinjiang Province, China | Lab stock |
| *Halomonas* LY02 | Wild-type *Halomonas* LY02 strain isolated from Turpan in Xinjiang Province, China | This study |
| *Halomonas* LY03 | Wild-type *Halomonas* LY03 strain isolated from Turpan in Xinjiang Province, China | This study |
| LYM1 | Recombinant LY03 with deleted *phaP*_1_ | This study |
| LYM1Δ*dddAC* | Recombinant LYM1 with deleted *dddAC* | This study |
| LYX | Recombinant LYM1 harboring *xylA-xfp* module integrated on IR_3724_ under control of P_porin59_ | This study |
| LYC | Recombinant LYM1 harboring *phaCAB* module integrated on IR_3724_ under control of P_J23100_ | This study |
| LYM2 | Recombinant LYM1 harboring ‘attP_int5_’ and ‘attP_bxb1_’ integrated on IR_0815_ and IR_3304_, respectively | This study |
| LYM3 | Recombinant LYM1 with ‘attP_bxb1_’ and ‘attP_int5_’ integrated on IR_3724_ and IR_1803_, respectively | This study |
| LYM4 | Recombinant LYM1 with ‘attP_bxb1_’ and ‘attP_int5_’ integrated on IR_2236_ and IR_1803_, respectively | This study |
| LYB | LYM1 derivate containing P_porin59_-*ydcW_Kp_-dhaT_Pp_* and P_porin51_-*orfZ* modules integrated on IR_3724_ and IR_1803_, respectively | This study |
| LYG | LYM1 derivate containing P_porin59_-*4hbd-sucD-ogdA* and P_porin51_-*orfZ* modules integrated on IR_2236_ and IR_1803_, respectively | This study |

### Table S2. Plasmids used in this study.

| **Plasmids** | **Description** | **Reference** |
| --- | --- | --- |
| pSEVA321 | Expression vector constructed with RK2 origin of replication and chloramphenicol resistance, Cm^R^ | ^4^ |
| pSEVA341 | Expression vector constructed with ColE1 origin of replication, kanamycin (Km^R^) and spectinomycin (Sp^R^) resistance | ^4^ |
| pRE112 | A suicide plasmid for gene knockout with R6K origin, Cm^R^, and 6×I-SceI sites | ^5^ |
| p321-P_J23114/105/100/111_-sfGFP | pSEVA321 derivates harboring *sfgfp* controlled by P_J23XXX_ promoters and RBS2000 | This study |
| p321-P_porin194/51/58/59/226/140/141_-sfGFP | pSEVA321 derivates harboring *sfgfp* controlled by P_porin_ promoter mutants and RBS2000 | This study |
| p321-*cinR*-P_Cin_- sfGFP | Inducible system constructed on pSEVA321, *cinR* controlled by P_LacI_, *sfgfp* controlled by P_Cin_ | This study |
| p321-*cymR*-P_Cym_-sfGFP | Inducible system constructed on pSEVA321, *cymR* controlled by P_LacIQ_, *sfgfp* controlled by P_Cym_ | This study |
| p341-*cymR*-P_Cym_-sfGFP | Inducible system constructed on pSEVA321, *cymR* controlled by P_LacIQ_, *sfgfp* controlled by P_Cym_ | This study |
| p321-*vanR*-P_Van_- sfGFP | Inducible system constructed on pSEVA321, *vanR* controlled by P_LacIQ_, *sfgfp* controlled by P_Van_ | This study |
| p321-*RNAP*-P_Mmp1_-sfGFP | Inducible T7-like system constructed on pSEVA321，MmP1 RNA polymerase controlled by P_LacIQ_, *sfgfp* controlled by P_MmP1_ | This study |
| pRE112-phaP_1_ | pRE112 derivate used for phaP_1_ knock-out. | This study |
| p341-P_Cym_-*dhaB-gdrAB* | pSEVA341 derivate containing *dhaB-gdrAB* (*K. pneumoniae*) expression module controlled by Cuma-inducible system | This study |
| p321-P_Van_-*pduP*_LR/ST_ | pSEVA321 derivate containing *pduP* (*L. reuteri* or *S. typhimurium*) expression module controlled by Van-inducible system | This study |
| p321-P_Cym_-*phaC-* P_Van_*-phaJ* | pSEVA321 derivate containing different *phaC* genes, including the ones from *P. entomophila* LAC32, *R. eutropha*, *A. hydrophila* 4AK4, and a chimeric *phaC*_AR_, controlled by Cuma-inducible system, together with *phaJ* from *P. entomophila* LAC32 and *A. hydrophila* 4AK4 controlled by Van-inducible system | This study |
| p321-J23110-sfGFP (TAAN/TGAN/TAGN) | pSEVA321 derivates harboring P_J23110_-sfGFP terminated by quadruplex stop codons TAAN/TGAN/TAGN | This study |
| p321-J23110-sfGFP-G4S-mRFP | pSEVA321 derivative harboring sfGFP (without a stop codon) and directly fused to mRFP (terminated by TAA stop codon) via a GGGGS linker under the control of P_J23110_ promoter and. | This study |
| p321-J23110-sfGFP-TAA-G4S-mRFP | pSEVA321 derivative harboring sfGFP (terminated by TAA) and then fused to mRFP (terminated by TAA) via a GGGGS linker under the control of P_J23110_ promoter. | This study |
| pRE112-IR_0815_-attP_int5_ | pRE112 derivate used to integrate attP_int5_ at loci IR_0815_, 1000 bp donor. | This study |
| pRE112- IR_0815_-attP_bxb1_ | pRE112 derivate used to integrate attP_bxb1_ at loci IR_0815_, 1000 bp donor. | This study |
| pRE112- IR_0815_-attP_int12_ | pRE112 derivate used to integrate attP_int5_ at loci IR_0815_, 1000 bp donor. | This study |
| pRE112- IR_3304_-attP_bxb1_ | pRE112 derivate used to integrate attP_bxb1_ at loci IR_3304_, 1000 bp donor. | This study |
| p321-Bxb1-P_J23110_-sfGFP | pSEVA321 derivate used to integrate sfGFP module mediated by Bxb1 integrase | This study |
| p321-Int12-P_J23110_-sfGFP | pSEVA321 derivate used to integrate sfGFP module mediated by Int12 integrase | This study |
| p321-Int5-P_J23110_-sfGFP | pSEVA321 derivate used to integrate sfGFP module (1 kb) mediated by Int5 integrase | This study |
| p321-Int5-*aldD_Pp_*-*dhaT_Pp_* | pSEVA321 derivate used to integrate *aldD_Pp_*-*dhaT_Pp_* (3 kb) by Int5 integrase | This study |
| p321-Int5-*4hbd-sucD-ogdA* | pSEVA321 derivate used to integrate *4hbd-sucD-ogdA* (5 kb) by Int5 integrase | This study |
| p321-Int5-*4hbd-sucD-ogdA-phaCAB* | pSEVA321 derivate used to integrate *4hbd-sucD-ogdA-phaCAB* (9 kb) by Int5 integrase | This study |
| psCre | pSEVA341 derivate used to excises the non-terget qesuence, including integrase gene and antibiotic marker, flanked by loxP sites mediated by sCre recombinase. | This study |
| pRE112-IRxxxx-attP_bxb1_ | pRE112 derivate used to integrate attP_bxb1_ at HENISs identified by SiteSeek, 1000 bp donor | This study |
| pRE112-IR_1803_-attP_int5_ | pRE112 derivate used to integrate attP_int5_ at IR_1803_ identified by SiteSeek, 1000 bp donor | This study |
| p321-P_Mmp1_-*xylA-xfp* | pSEVA321 derivate harboring *xylA-xfp* controlled by P_Mmp1_ induced by IPTG | This study |
| p321-P_porin58/59/226/140_-*xylA-xfp* | pSEVA321 derivates harboring *xylA-xfp* controlled by P_porin58/59/226/140_ | This study |
| p321-Int5-P_porin59_-*xylA-xfp* | pSEVA321 derivate used to integrate *xylA-xfp* module using Int5 integrase | This study |
| p321-Int5-P_J23100_-*phaCAB* | pSEVA321 derivate used to integrate *phaCAB* module using Int5 integrase | This study |
| p321-P_porin58_-*aldD_Pp_*-*dhaT_Pp_*-P_J23111_-*orfZ* | pSEVA321 derivate containing *aldD*-*dhaT* cluster from *Pseudomonas putida* KT2440 controlled by P_porin58_, and *orfZ* from *Clostridium kluyveri* controlled by P_porin194_ | This study |
| p321-P_porin58_-*aldD_Pp_/aldD_Hb_/aldH*_Ec_*/gabD4*_Re_*/aldD*_Kp_-  *dhaT_Pp_/adhP_Hb_-*P_porin194_-*orfZ* | pSEVA321 derivates harboring genes encoding aldehyde dehydrogenase and alcohol dehydrogenase from different hosts controlled by P_porin58_ and *orfZ* controlled by P_porin194_ | This study |
| p321-Bxb1-P_porin59_-*aldD_Kp_*-*dhaT_Pp_* | pSEVA321 derivate used to integrate *aldD*_Kp_-*dhaT_Pp_* module using Bxb1 integrase | This study |
| p321-Bxb1-P_porin194_-*4hbd-sucD-ogdA* | pSEVA321 derivate used to integrate *4hbd-sucD-ogdA* using Bxb1 integrase | This study |
| p321-Int5-P_porin51_-*orfZ* | pSEVA321 derivate used to integrate *orfZ* module using Int5 integrase | This study |

### Table S3. Brief summary of whole-genome sequencing of *Halomonas* LY03.

|  |  | Category | Information (numbers) |
| --- | --- | --- | --- |
| Genome | DNA sequencing | Genome size (bp) | 4,141,019 |
|  | Annotation | Total annotations | 4,220 |
|  |  | CDS | 3,904 |
|  |  | tRNA | 62 |
|  |  | rRNA | 18 |
|  |  | sRNA | 8 |
|  |  | CRISPR | 1 |
|  |  | Tandem repeat | 227 |

### Table S4. Comparison of the molecular weight of PHB synthesized by LY03, LYM1, and TDH4.

| **Strain** | **M_P_（Da）** | **M_n_（Da）** | **M_w_（Da）** | **M_z_（Da）** |
| --- | --- | --- | --- | --- |
| LY03 | 635,209 | 482,284 | 761,968 | 1,199,668 |
| LYM1 | 495,489 | 314,258 | 553,922 | 965,191 |
| TDH4^a^ | 411,379 | 310,486 | 479,614 | 746,475 |

* The data are all generated from this study. M_P_: peak molecular weight (the molecular weight at the peak of the distribution curve); M_n_: number-average molecular weight, $M_{n}=\sum N_{i}M_{i}/\sum N_{i}$; Mw: weight-average molecular weight, $M_{w}=\sum N_{i}M_{i}^{2}/\sum N_{i}M_{i}$; Mz: Z-average molecular weight, $M_{z}=\sum N_{i}M_{i}^{3}/\sum N_{i}M_{i}^{2}$. $N_{i}$ and $M_{i}$ are the number and molecular weight of species $i$.

^a^ TD01-derived strain with defected *phaP* constructed by Shen et al. donated from THU group^3^.

### Table S5. Constitutive promoters of P_J23XXX_ and P_porin_ used in this study.

| **Promoter** | **Sequence** | **Source** |
| --- | --- | --- |
| P_J23114_ | TTTATGGCTAGCTCAGTCCTAGGTACAATGCTAGC | iGEM BBa_J23114 |
| P_J23105_ | TTTACGGCTAGCTCAGTCCTAGGTACTATGCTAGC | iGEM BBa_J23105 |
| P_J23100_ | TTGACGGCTAGCTCAGTCCTAGGTACAGTGCTAGC | iGEM BBa_J23105 |
| P_J23111_ | TTGACGGCTAGCTCAGTCCTAGGTATAGTGCTAGC | iGEM BBa_J23111 |
| P_porin194_ | TTGCGTTCACTGGAATCCCAGTATCTAATTTGACCTGCGAGCA | ^3^ |
| P_porin51_ | TTGCGTTCACTGGAATCCCATTTTAGAGTTTGACCTGCGAGCA | ^3^ |
| P_porin58_ | TTGCGTTCACTGGAATCCCAATATAGAGTTTGACCTGCGAGCA | ^3^ |
| P_porin59_ | TTGCGTTCACTGGAATCCCAGTATAGAGTTTGACCTGCGAGCA | ^3^ |
| P_porin226_ | TTGCGTTCACTGGAATCCCAGTATAAAGTTTGACCTGCGAGCA | ^3^ |
| P_porin140_ | TTGCGTTCACTGGAATCCCAGTATAAGATTTGACCTGCGAGCA | ^3^ |
| P_porin141_ | TTGCGTTCACTGGAATCCCAGTATATACTTTGACCTGCGAGCA | ^3^ |

### Table S6. Composition and content of dominant fatty acids in the waste soybean oil from ‌Foshan Haitian Flavouring And Food Co., Ltd.

| **Component** | **Content (wt%)** |
| --- | --- |
| C8:0 (Octanoic acid) | ND |
| C10:0 (Decanoic acid) | ND |
| C12:0 (Dodecanoic acid) | ND |
| C14:0 (Myristic acid) | ND |
| C16:0 (Palmitic acid) | 11.9 |
| C18:0 (Stearic acid) | 4.1 |
| C18:1 (Oleic acid) | 17.7 |
| C18:2 (Linoleic acid) | 55.3 |
| C18:3 (α-Linolenic acid) | 9~9.5 |

ND, not detected. The residual ratio not included in the table indicates water content (1.5~2**%**).

### Table S7. BLAST analysis of PduP (from *Lactobacillus reuteri* ) in *Halomonas* LY03.

| **Enzyme name** | **Percentage identity*** | **E value*** | **Query cover*** |
| --- | --- | --- | --- |
| LY03GL003351 | 23.02% | 2e-05 | 74% |
| LY03GL002807 | 21.39% | 0.037 | 35% |

* The data of percentage identity (per. ident), E-value, and query coverage were obtained from the NCBI database.

### Table S8. Percentages of TAA, TGA and TAG stop codons in the gene sets grouped by protein abundance, mRNA abundance and protein-to-mRNA ratio (translation efficiency) from top 100% to top 5% (1 g L⁻¹ urea).

|  | Protein/mRNA | Protein | mRNA |
| --- | --- | --- | --- |
| TAA | 51.2%→68.0%⬆ | 51.2→82.5%⬆ | 51.2→66.0%⬆ |
| TGA | 29.3%→22.0%⬇ | 29.3→12.0%⬇ | 29.3→19.5%⬇ |
| TAG | 19.5%→10.0%⬇ | 20.5→5.5%⬇ | 20.5→14.5%⬇ |

### Table S9. Percentages of TAA, TGA and TAG stop codons in the gene sets grouped by protein abundance, mRNA abundance and protein-to-mRNA ratio (translation efficiency) from top 100% to top 5% (5 g L⁻¹ urea).

|  | Protein/mRNA | Protein | mRNA |
| --- | --- | --- | --- |
| TAA | 51.2%→65.5%⬆ | 51.2→82.0%⬆ | 51.2→67.0%⬆ |
| TGA | 29.3%→24.5%⬇ | 29.3→11.5%⬇ | 29.3→19.0%⬇ |
| TAG | 19.5%→10.0%⬇ | 20.5→6.5%⬇ | 20.5→14.0%⬇ |

### Table S10. Parameters, descriptions, and recommended setting values for SiteSeek

| **Parameter** | **Description** | **Recommended Value** |
| --- | --- | --- |
| target_length | Recommanded length of selected regions for preliminary HENIS screening | 350 |
| min_tf_len | Minimum sequence length required for TFBs alignment within the ‘target_length’ | 8 |
| buffer | Buffer size flanking the excluded region to ensure the following effective filtering | 10 |
| similarity | Identital threshold of TFBs aligned to the ‘tf_len’ fragments. The sequences with ‘similarity’ exceeding the recommended value are removed from the ‘target_length’ for next round analysis. | 0.9 |
| Threshold | Threshold for promoter prediction using ‘BDGP’ toolkit | 0.9 |
| min_segment | Minimum aequence length of HENIS region after removing the TFB and promoter element | 100-145 |
| min_taa_len | Minimum distance of HENIS region from adjacent translation termination position | 150 |
| Intensity | Omics filtering index for HENIS site screening, applied when transcriptomic FPKM ranks of flanking genes exceed the specified percentile level (0-1, or 0-100%) | 0.1 |
| Number_of_sites | Expected number in total for the identified HENIS candidates | 10~20 |

### Table S11. Parameters used for HENIS screening in *Halomonas* LY03 and LY01 using Siteseek.

| **LY03** | | **LY01** | |
| --- | --- | --- | --- |
| **Filter criteria** | **IR number** | **Filter criteria** | **IR number** |
| IR*_i_* length ≥ 350 bp | 205 | IR*_i_* length ≥ 350 bp | 206 |
| IR without tRNA, rRNA, sRNA, TRF | 148 | IR without tRNA, rRNA, sRNA, TRF | 145 |
| IR without TF binding site (similarity = 0.9) | 130 | IR without TF binding site (similarity = 0.9） | 125 |
| IR*_i_* length ≥ 145 bp after potential promoter and terminator removal | 44 | IR*_i_* length ≥ 100 bp after potential promoter and terminator removal | 48 |
| RNA: Gene*_x&x+1_* in the top 10%  Number_of_sites = 10 | 10 | RNA: Gene*_x&x+1_* in the top 10%  Number_of_sites = 3 | 3 |

**Table S12. High-expression neutral integration site (HENIS, letters in bold) identified by SiteSeek.**

| Host Loci name | | Segment length includes loci (bp) | Inserted position  (bp, 5’- to 3’-end) |
| --- | --- | --- | --- |
| LYM1 | IR_1803_ | 309 | 108 |
|  | IR_2236_ | 294 | 203 |
|  | IR_2560_ | 223 | 220 |
|  | IR_2720_ | 153 | 127 |
|  | IR_2840_ | 251 | 187 |
|  | IR_2846_ | 264 | 167 |
|  | IR_3192_ | 332 | 316 |
|  | IR_3385_ | 213 | 126 |
|  | IR_3507_ | 152 | 76 |
|  | IR_3724_ | 177 | 130 |
|  | IR_0968_ | 110 | 50 |
| LY01 | IR_1485_ | 215 | 150 |
|  | IR_3495_ | 241 | 170 |

Note: IR_1803_ denotes an insertion region located in the intergenic region between genes numbered LY03001803 and LY03001804. The failures of integrated loci are marked with gray background.

**Table S13. Molecular weight (Mw) of PHB produced by two-stage continuous and normal fed-batch processes at 20-m^3^ scale**.

| Harvest cycles / Batch | Two-stage continuous | Normal fed-batch |
| --- | --- | --- |
| 1 | 958,715 | 408,752 |
| 2 | 954,102 | 760,146 |
| 3 | 816,610 | 659,917 |
| 4 | 910,542 | 533,219 |
| 5 | 660,978 | / |
| 6 | 868,795 | / |
| 7 | 654,954 | / |
| Mean ± SD | 832,099 ± 131,472 | 590,509 ± 149,784 |
| CV (%) | 15.8 | 25.4 |

* The data are all generated from this study. CV = (|SD|/Mean) × 100%.

**Supplementary Figures**

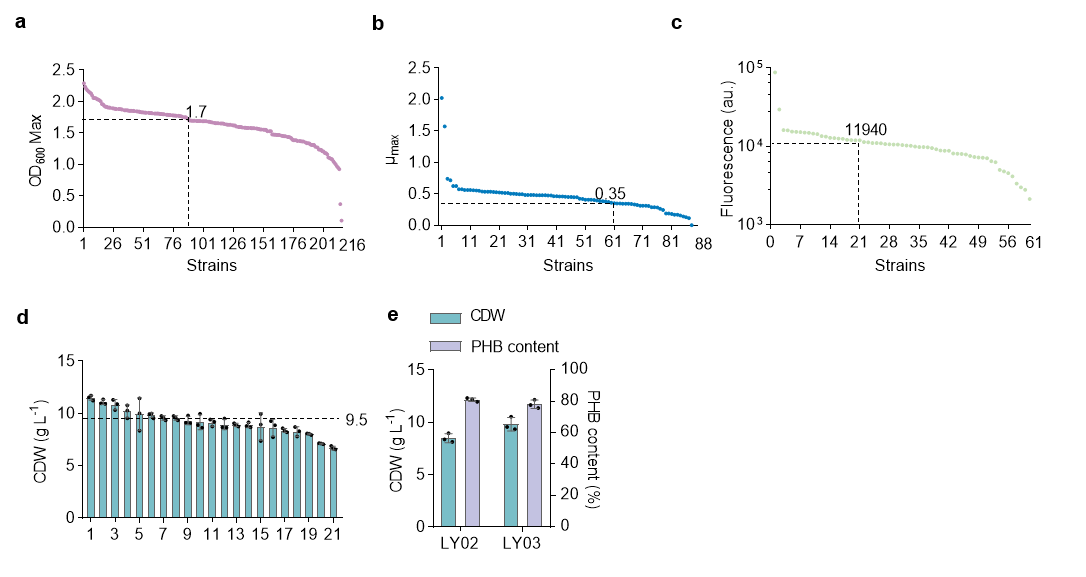

### Figure S1. Details of the screening process for high-performing halophilic isolates.

(**a**) OD_600_ of isolated strains cultured in a 96-well plate containing 50MMG medium for 24 h. 88 strains with OD_600_ over 1.7 were selected for specific growth analysis in part **b**. (**b**) Maximum specific growth rate (µ_max_) of each strain from part **a** grown in a 96-well plate for 24 h (50MMG). 61 Strains with μ_max_ over 0.35 were selected for high-through PHB accumulation assay in part **c**. (**c**) Fluorescence intensity (FI), determined by FACS (flow cytometer), of BODIPY-stained cells by isolated strains in part **b** after 48-h growth in 50MMG. 21 Strains with FI > 11,940 were selected for shake flask study of CDW and PHB accumulation in part **d**. (**d**) Cell growth assessment of the selected 21 strains cultuered in a 150-mL shake flask containing 20 mL 50MMG medium. 8 Strains with CDW higher than 9.5 g L^-1^ were selected for 16S rRNA sequencing. (**e**) Shake flask study of CDW and PHB content by two isolated halophilic strains, *Halomonas* LY02 and LY03, of distinct 16S rRNA sequence. Error bars in **d** and **e** represent standard deviations, n = 3. For 96-well plate study in **a-c**, n = 1.

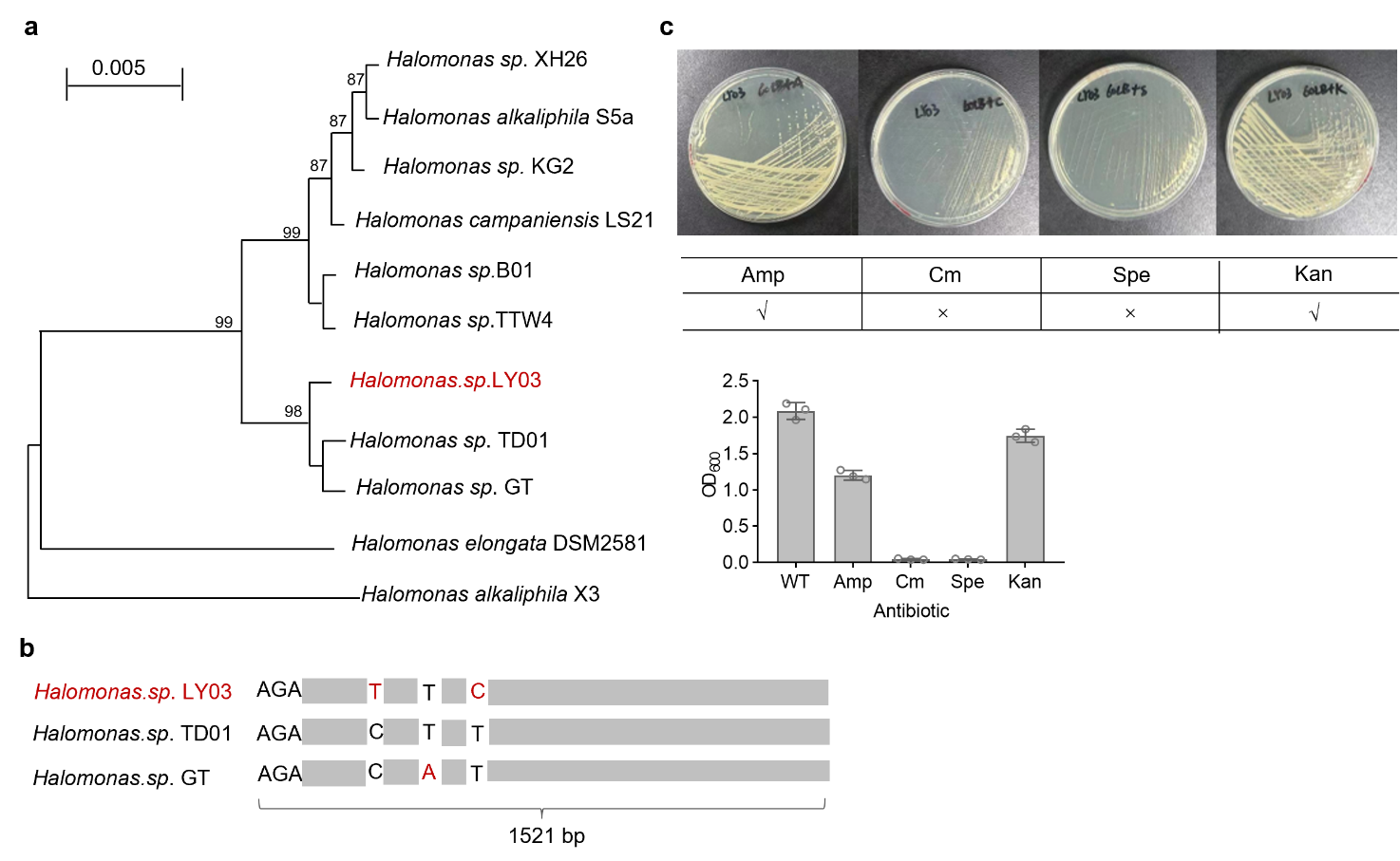

### Figure S2. Phylogenetic analysis and antibiotic resistance testing of *Halomonas* LY03.

(**a**) Phylogenetic tree of *Halomonas* LY03 against different halophiles based on 16S rRNA sequence using MEGA 11.0. (**b**) BLAST alignment of 16S rRNA genes from three *Halomonas* strains, LY03, TD01^6^, and GT^7^. Grey blocks indicate aligned regions, and variant nucleotides are indicated at mutation sites. (**c**) Antibiotic resistance testing of *Halomonas* LY03 grown on an agar plate containing different different antibiotics for 12 h (upper panel), and a 150-mL shake flask (OD_600_ values) containing 20-mL 60LB medium supplemented relevant antibiotics for OD_600_ measurement after 24-h incubation. For shake flask study, error bars in **c** represent standard deviations, n = 3.

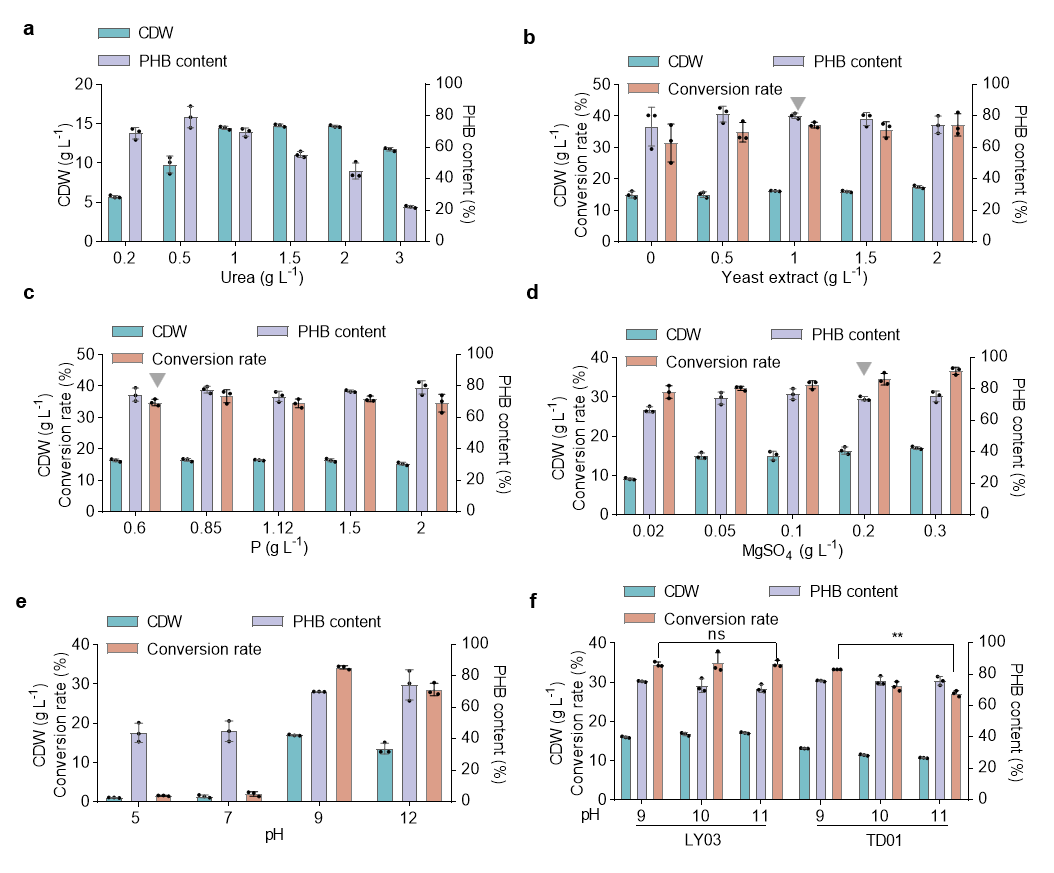

### Figure S3. Medium optimization and alkali-tolerance assessment of *Halomonas* LY03.

(**a**) CDW and PHB content of *Halomonas* LY03 grown in 50MMG medium of different C/N ratio (30 g L^-1^ glucose + 0.2, 0.5, 1, 1.5, 2, and 3 g L^-1^ urea). (**b**-**d**) CDW and PHB content of *Halomonas* LY03 grown in a 150-mL shake flask containing 20-mL 50MMG medium with optimized C/N ratio (35 g L^-1^ glucose + 1 g L^-1^ urea) supplemented with different concentration of yeast extract (**b**), phosphate (**c**), and magnesium sulfate (**d**), respectively. (**e**) pH-tolerance assessment of *Halomonas* LY03 grown in optimized 50MMG medium with pH adjusted to 5, 7, 9, and 12, respectively. (**f**) Comparison of CDW and PHB content by *Halomonas* LY03 grown on different pH conditions in optimized 50MMG medium compared to *Halomonas* TD01^8^ grown in the same pH coditions. Error bars represent standard deviations, n = 3.

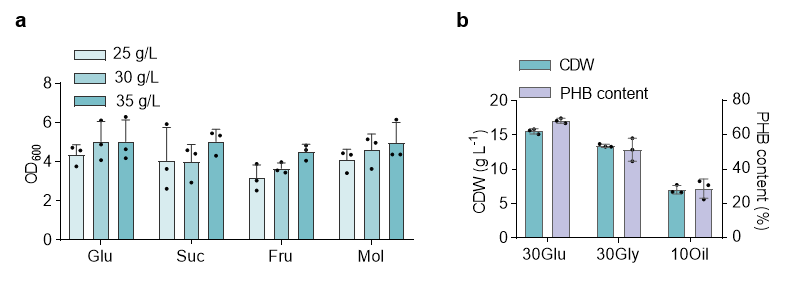

### Figure S4. Assessing the carbon source-utilization capability of *Halomonas* LY03.

(**a**) OD_600_ of *Halomonas* LY03 grown on different carbon sources, including glucose (Glu), sucrose (Suc), fructose (Fru), and molasses (Mol), in a 96-well plate for 24 h. Specifically, 25, 30, and 35 g L^-1^ of each sugar were respectively supplemented for growth analysis (OD_600_). (**b**) Shake flask study of CDW and PHB content by *Halomonas* LY03 grown in a 150-mL shake flask containing 50MM medium supplemented with 30 g L^-1^ glucose and glycerol, and 10 g L^-1^ waste soybean oil, respectively. Error bars represent standard deviations, n = 3.

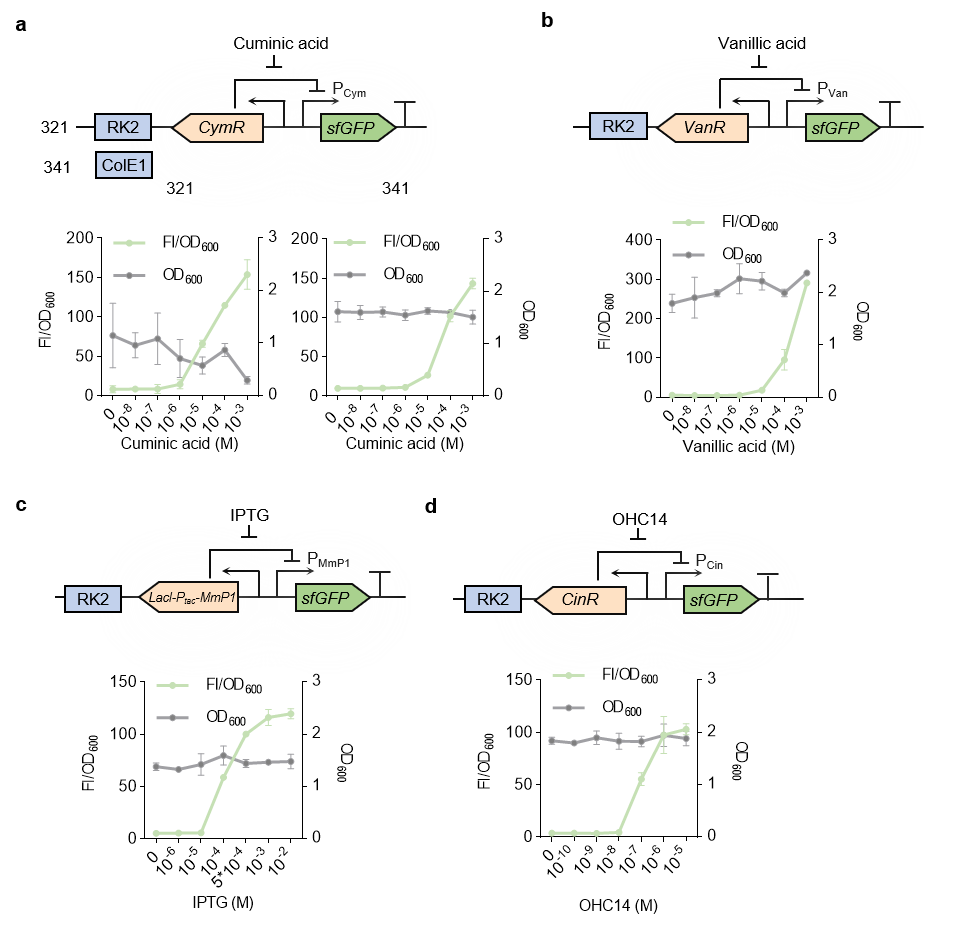

### Figure S5. Dose-response characterization of different inducible systems in *Halomonas* LYM1.

(**a**-**d**) Dose-response curve chracterization of different inducible systems, indlucing cuminic acid- (constructed in vectors with high- (colE1) and medium-copy (RK2) origin), Vanillic acid-, IPTG-, and OHC14-induced system, by recombinant LYM1 using sfGFP as a reporter. Fluorescent intensity of each cell culture was normalized by OD_600_ (FI/ OD_600_). Error bars represent standard deviations, n = 3.

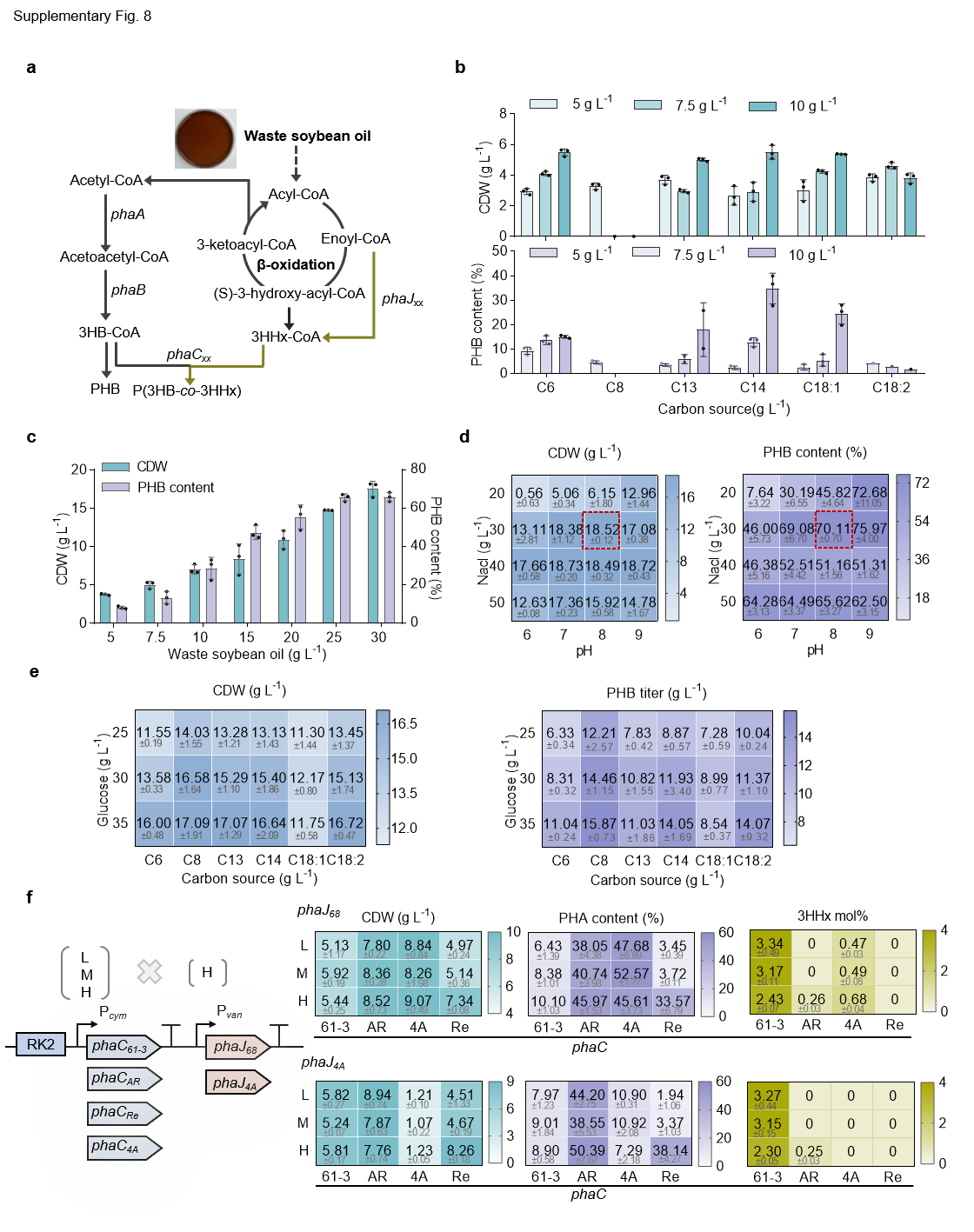

### Figure S6. Engineering PHB and PHBHHx production by recombinant LYM1 from fatty acids.

(**a**) Metabolic pathways for PHB and PHBHHx synthesis from waste soybean oil mainly containg different fatty acids in recombinant *Halomonas* LYM1. (**b**) Shake flask study of CDW and PHB content by LYM1 grown on different fatty acids of various carbon-chain length. (**c**) Shake flask of CDW and PHB content by LYM1 grown in 50MM meiudm supplemented with different concentration of waste soybean oil. (**d**) Orthognal assessment of different NaCl concentration and pH level that affect the CDW and PHB content by LYM1 grown on 30 g L^-1^ waste soybean oil. (**e**) Assessing the co-utilization of glucose and different fatty acids (2 g L^-1^) by LYM1 for yielding PHB production. (**f**) Shake flask study of CDW and PHBHHx production by recombinant LYM1 carrying different *phaC-phaJ* cluster combinations grown on 30g L^-1^ waste soybean oil. Notes: *phaC*_61-3_ (61-3), *phaC*_Re_ (Re), *phaC*_4A_ (4A), and *phaC*_AR_ (AR) indicate different PHA synthase from *Pseudomonas sp.* 61-3, *Ralstonia eutropha* H16, *A. hydrophila* 4AK4, and a chimeric enzyme composed of 26% of the N-terminal of *PhaC*_Ac_ from *Aeromonas caviae* and 74% of the C-terminal of *PhaC*_Re_, respectively; *phaJ*_68_ and *phaJ*_4A_ indicate (*R*)-specific enoyl-CoA hydratase from *Pseudomonas putida* 1668 and *A.shydrophila* 4AK4. Letters of L, M, and H indicate low-, meium- and high-expression level of *phaC* induced by 0.01, 0.1, and 1 mM cumaric acid, while *phaJ* was induced by 1 mM vanlinic acid of high expression level. Error bars represent standard deviations, n = 3; The ‘mean + SD’ values in heatmaps were obtained from triplicate samples.

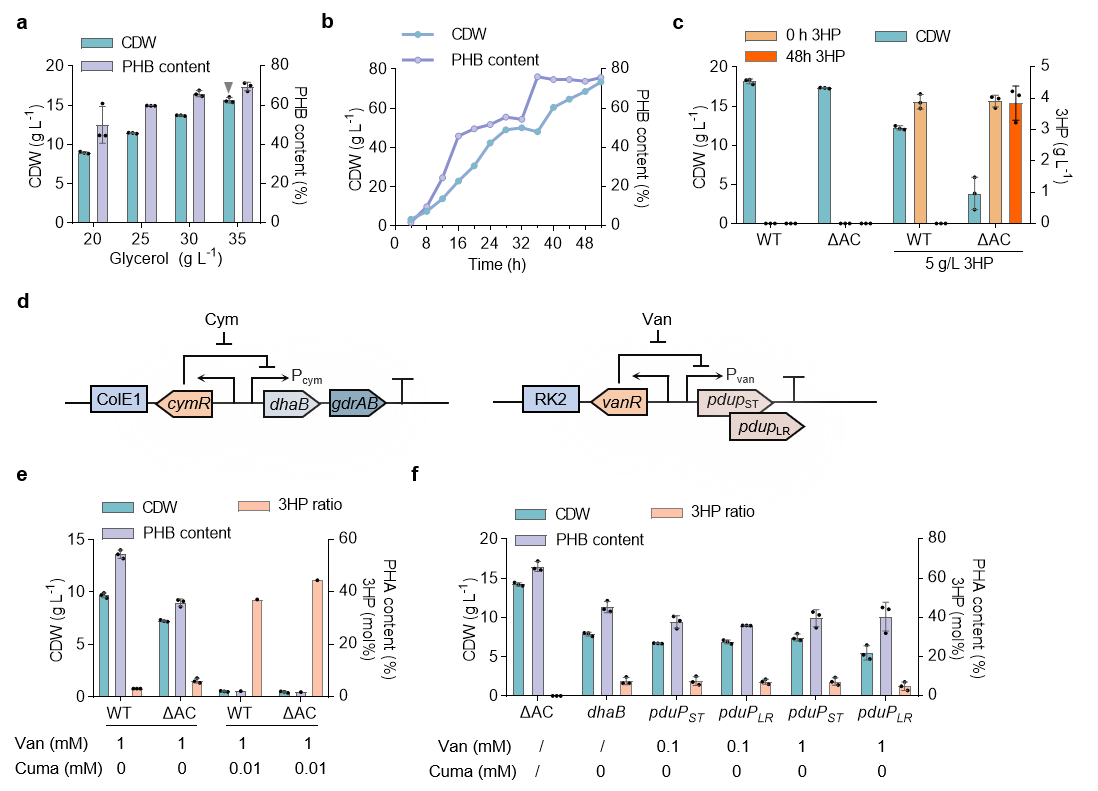

### Figure S7. Engineering PHB and PHBP production by *Halomonas* LYM1 from glycerol.

(**a-b**) Shake falsk (**a**) and fed-batch (**b**) study of CDW and PHB content by *Halomonas* LYM1 grown glycerol only. (**c**) Studying the effects of *dddAC* deletion on cell growth and 3-hydroxypropionate (3HP) metabolism in recombinant LYM1. (**d**) Schematic diagram of plasmids harboring two metabolic modules, *dhaB-gdrAB* driven by P*cym* and *pduP*_ST/LR_ driven by P*_van_*, which convert glycerol into 3HP-CoA, constructed on pSEVA341 and pSEVA321, respectively. (**e**) Shake flask study of CDW and poly(3-hydroxybutyrate-*co*-3-hydroxypropionate) (PHBP) by LYM1 (WT) and its derivative with defective *dddAC* genes (ΔAC) harboring 3HP-CoA synthesis modules using glycerol as a sole carbon source under the induction of vanlinic acid and cumaric acid. (**f**) Studying the effect of *pduP* gene from different host on CDW and PHBP accumulation by recombinant LYM1Δ*dddAC* (ΔAC) grown on 30 g L^-1^ crude glycerol only. For shake flask studies, error bars represent standard deviations, n = 3. For fed-batch study in **b**, n = 1.

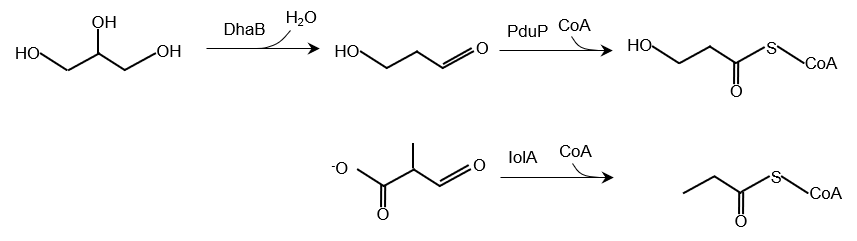

### Figure S8. Bioreactions catalyzed by and PduP-like enzymes in LY03.

Two predicted copies of *iolA* gene, including LY03GL003351 and LY03GL002807, encoding CoA-acylating methylmalonate-semialdehyde dehydrogenase, which show high identity to *pduP* gene encoding propanal dehydrogenase of oxidoreductase and Coenzyme A transferase function, are potentially to catalyze aldehyde group to acyl coenzyme A. The BLAST analysis of bioreactions was predicted using UniProt.

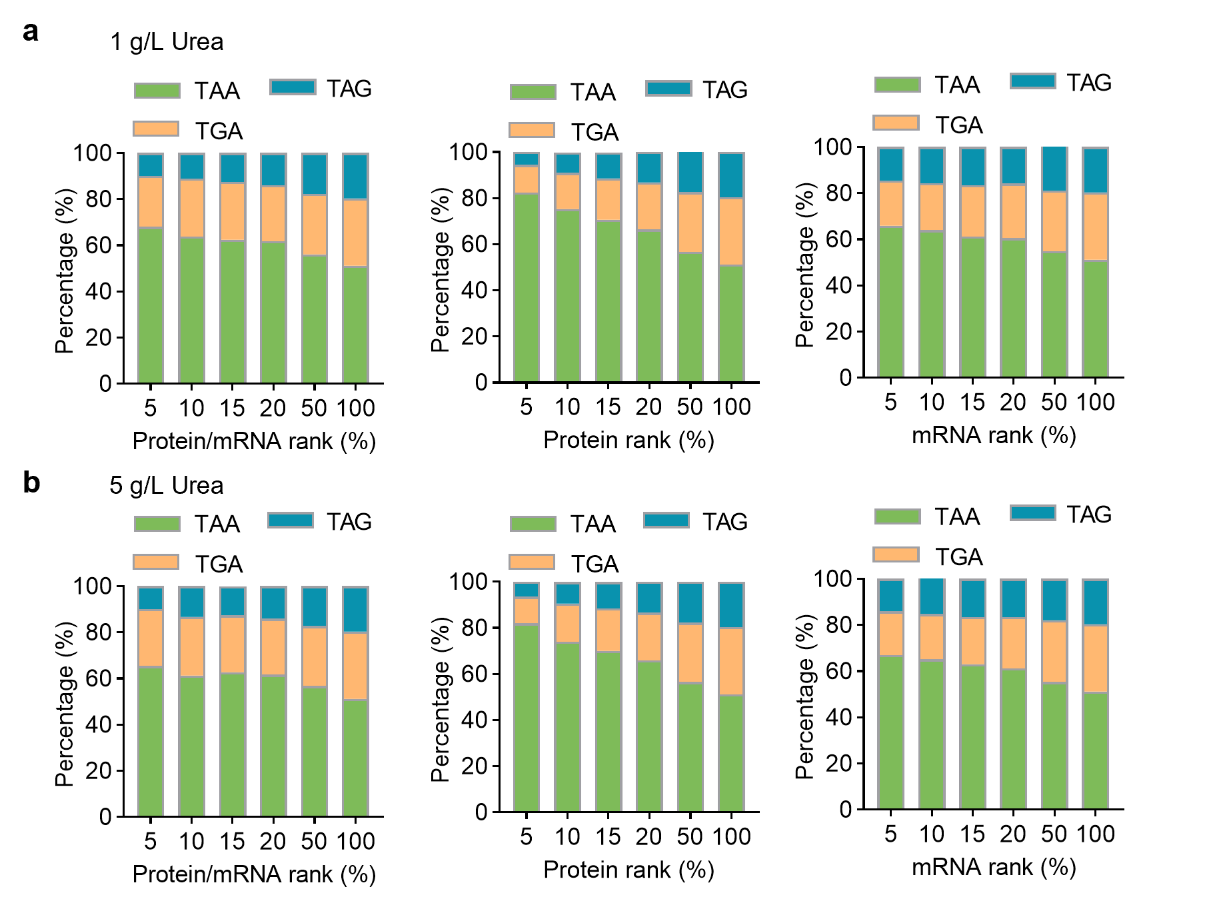

### Figure S9. Distribution of TAA, TAG, and TGA in gene sets grouped by varied top expression levels (top 5%, 10%, 15%, 20%, 50% and 100%), including translation efficiency, protein level, and mRNA level, by *Halomonas* LYM1 grown on 1 g L⁻¹ (a) and 5 g L⁻¹ (b) urea, respectively.

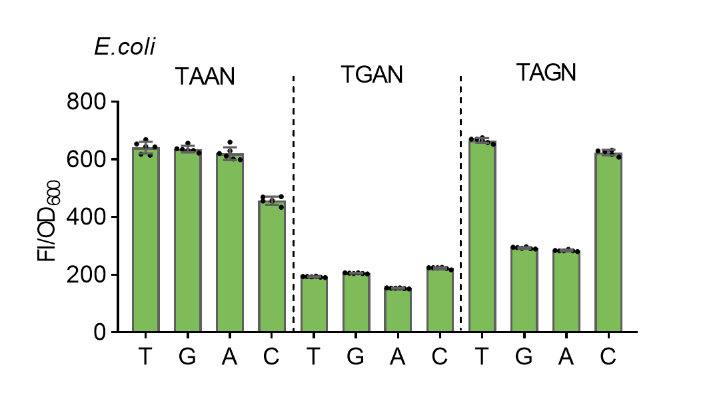

### Figure S10. Comparative analysis of normalized fluorescence in recombinant *E. coli* S17-1 harboring sfGFP reporter with terminated translation using quadruplex stop codon TAAN, TAGN, or TGAN.

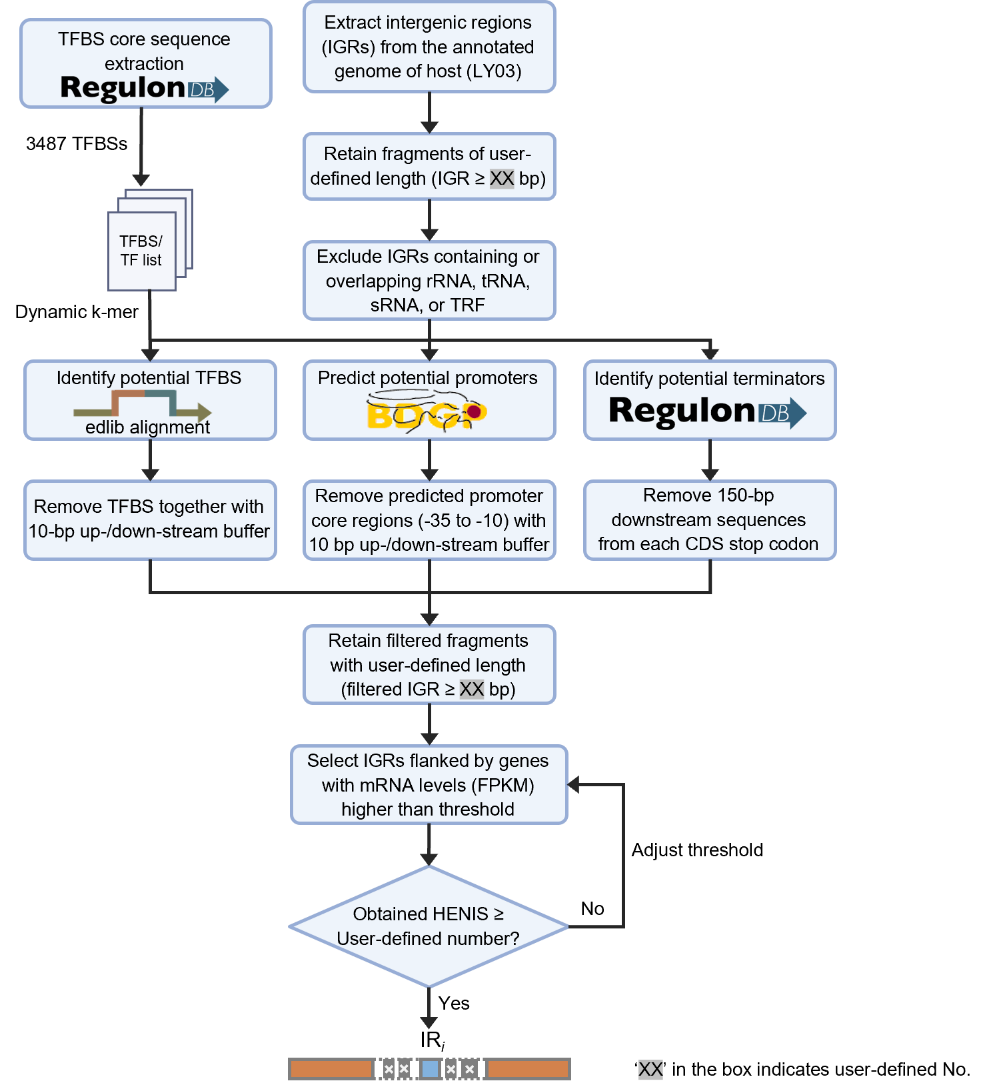

### Figure S11. The ‘SiteSeek’ workflow for screening high-expression neutral integration sites (HENISs).

**I**) The intergenic regions (IGRs) consisting of non-coding sequence were firstly generated by removing the annotated coding sequence (CDS); **II**) The IGRs with sequence length no shorter than the user-defined value were retained; **III**) The IGRs overlapping with (or containing) functional genomic elements including rRNAs, tRNAs, sRNAs, and tandem repeat fragments (TRFs) were excluded; **IV**) IGR candidates were then subjected to three filtering processes to eliminate potential interferences, including removal of transcription factor binding sites (TFBSs, namely operator core region), promoter-associated regions predicted by neural network-based promoter prediction^10^, and potential terminator regions (150 bp away from the stop codon of each CDS). The filtered ones with sequence length no shorter than 145 (user-defined parameter) bp were retained; **V**) The resultant IGRs were further analyzed using the user-input transcriptomic data (optional) to generate desired numbers of HENISs for genomic integration test. Detailed code for whole-process HENISs screening can be accessed in GitHub (https://github.com/Halomonas/SiteSeek).

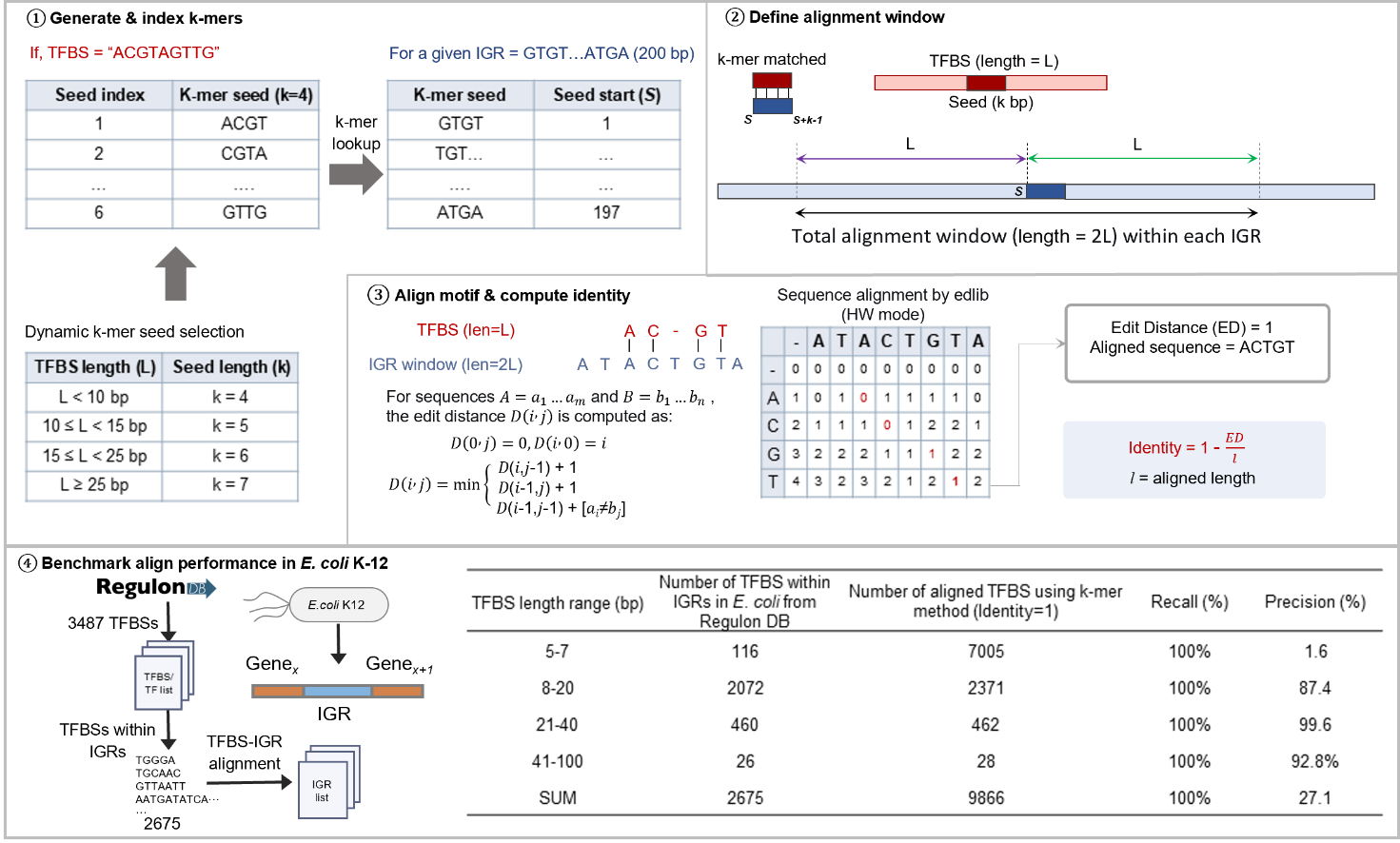

### Figure S12. TFBS alignment using an optimized dynamic k-mer algorithm.

Core TFBS motifs generated from ‘RegulonDB’ were decomposed into variable-length k-mer seeds according to the length of TFBSs for fast lookup against the k-mer seeds within each IGR to identify the k-mer matched regions. The k-mer matched regions were further extended into full alignment windows for semi-global alignment (edlib, HW mode), the edit distance and alignment length were used to calculate sequence identity. The recall and precision of this alignment strategy were validated by mapping the IGR-resident TFBS from ‘Regulon DB’ to the IGRs within the genome of *E. coli* K-12. Particularly, the length of core TFBS motif shorter than 8 bp significantly affects the alignment precision due to the high-frequency mis-match.

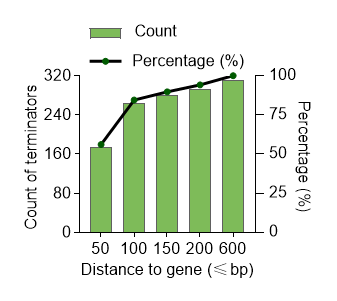

### Figure S13. Distance distribution of the experimentally characterized terminators to the upstream stop codon in *E. coli.*

More than 90% of terminators are located within 150 bp downstream of their corresponding genes (RegulonDB dataset)^9^.

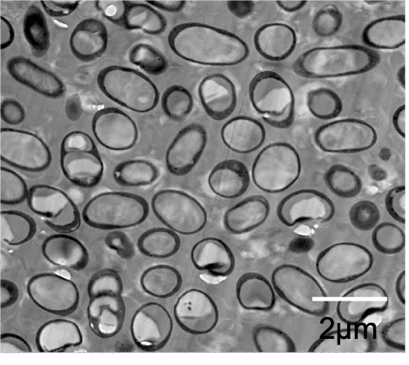

### Figure S14. Transmission electron micrograph (TEM) analysis of LY01 with natural single-granule PHB accumulation*.*

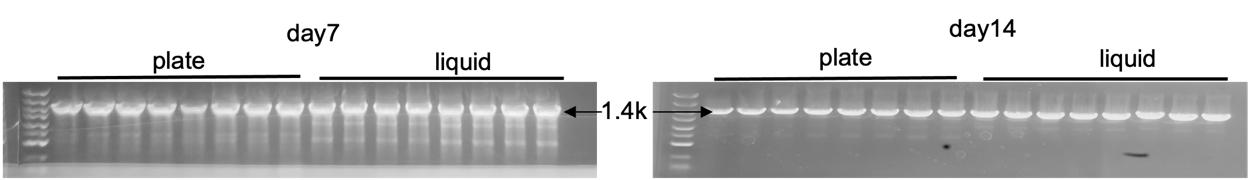

### Figure S15. Robustness validation of chromosomally engineered LYM1 strains harboring sfGFP expression module via colony PCR after 7- (left) and 14- (right) round day-by-day passaging growth on an agar plate (plate) and a shake flask (liquid).

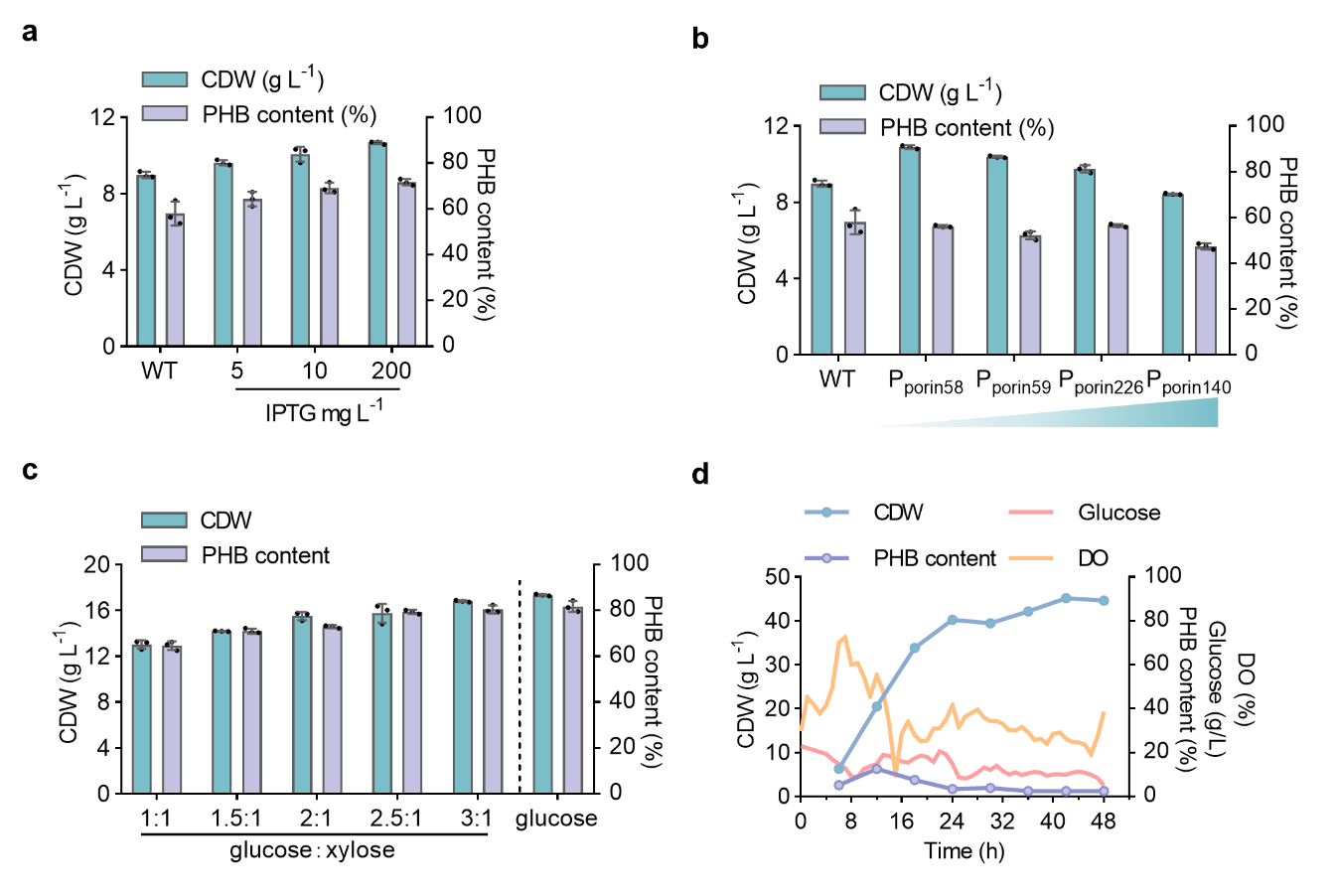

### Figure S16. Fine-tuning the chromosomal expression of *xylA-xfp* gene cluster in recombinant LYM1.

(**a**) Shake flask study of CDW and PHB content by recombinant LYM1 carrying p321-P_Mmp1_-*xylA-xfp* plasmid grown in 50MM medium containing 20 g L^-1^ glucose and 10 g L^-1^ xylose induced by different concentration of IPTG. (**b**) Shake flask study of CDW and PHB content by recombinant LYM1 carrying p321-P_porin lib_-*xylA-xfp* plasmids, the strength of selected P_porin_ promoters was of closely induction level of the optimal induction group from part **a**, grown in 50MM medium containing 20 g L^-1^ glucose and 10 g L^-1^ xylose. (**c**) Cell growth (CDW) and PHB content by recombinant LYX grown on mixed carbon source containing different glucose-to-xylose ratios (within 36 g L^-1^ sugar in total) in 50MM medium. **(d)** Time-course profiles of CDW, PHB content, dissolved oxygen (DO) and residual glucose by LYM1 grown in a 5-L bioreactor using glucose and xylose ((3:1, w/w)) as a feedstock. Error bars represent standard deviations, n = 3. For fed-batch study, n = 1.

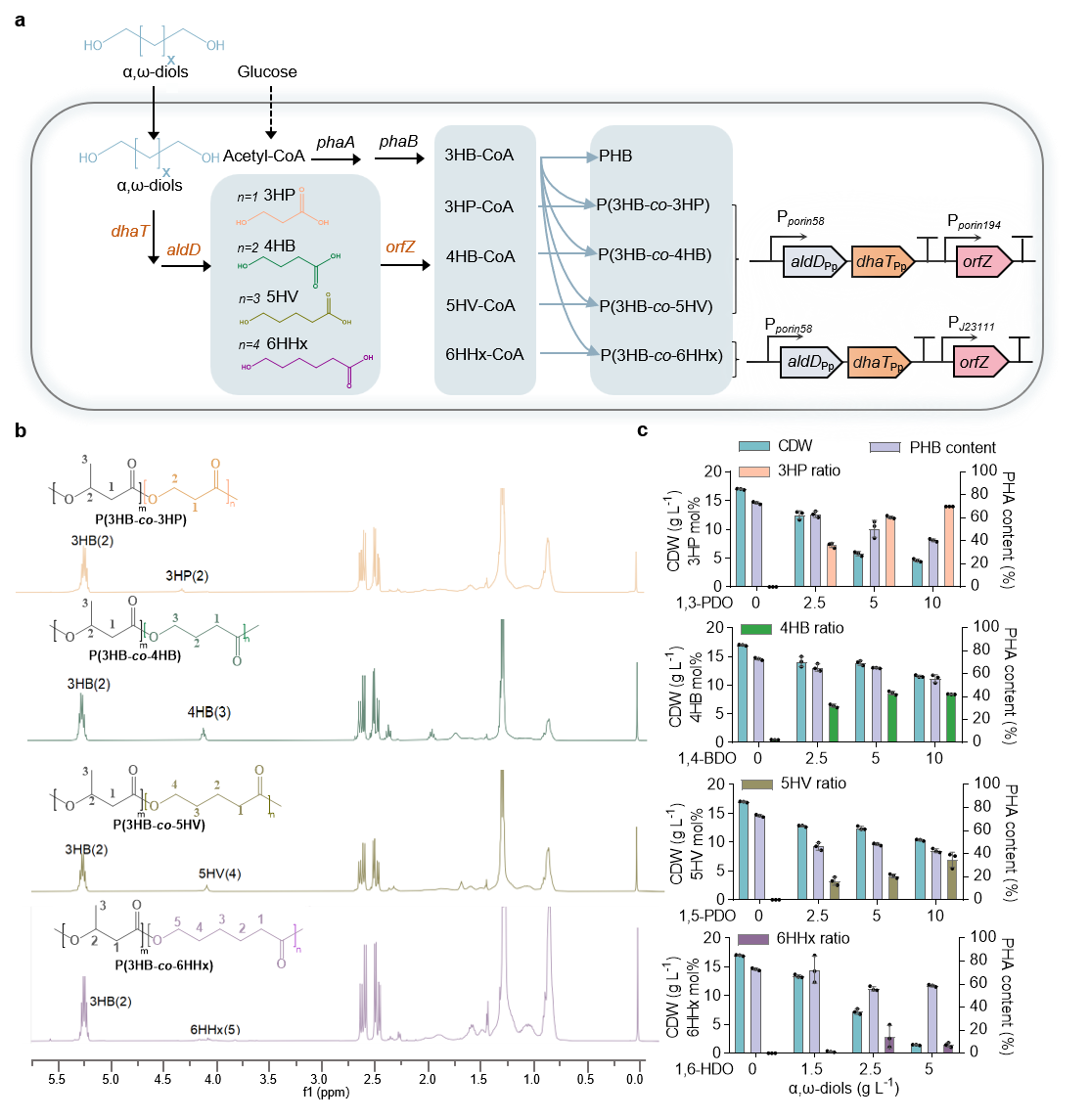

### Figure S17. Biosynthesis of PHA copolymers from glucose and α,ω-diols by recombinant LYM1.

(**a**) Metabolic pathway diagram for different PHA copolymers synthesis, including P(3HB-*co*-3HP), P(3HB-*co*-4HB), P(3HB-*co*-5HV), and P(3HB-*co*-6HHx), from glucose and α, ω-diols (C3 to C6) by recombinant LYM1. (**b**) ^1^H NMR spectra of different obtained PHA copolymers. The characteristic peaks of 3HB and different α,ω-hydroxyalkonic monomers are highlighted. (**c**) Shake flask study of CDW, PHA content, and α,ω-monomer ratio by recombinant LYM1 grown in 50MMG medium supplemented with different concentration of relevant α,ω-diols. Error bars represent standard deviations, n = 3.

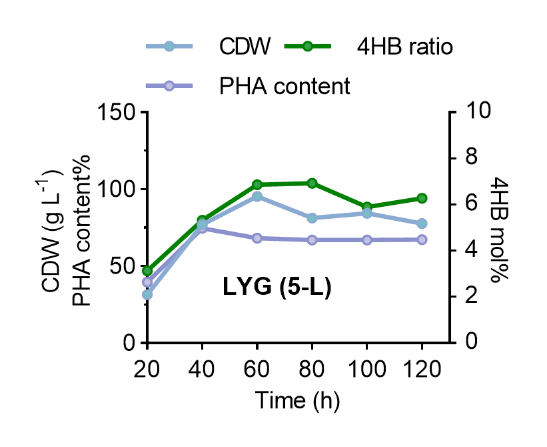

### Figure S18. Two-stage continuous fermentation of P34HB by LYG conducted in a 5-L bioreactor using glucose as a sole carbon source.

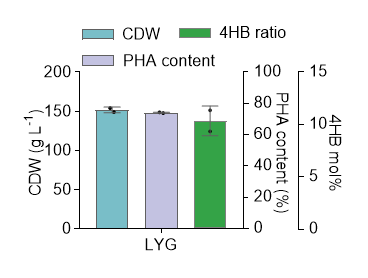

### Figure S19. High cell-density fed-batch fermentation of chromosomally engineered LYM1 harboring 4HB-synthesis module with higher expression level at 20 m³ scale using custom-optimized fermentation process.

Error bars represent standard deviations, n = 2.

### Supplementary Note. Calculation of production-efficiency-per-volume ratio (*P*_v_) of two-stage continuous against fed-batch study.

The function of production-efficiency-per-volume ratio (*P*_v_) is defined as:

*P_v_*$=\frac{n*\frac{V*\alpha}{V_{c}^{total}}}{2*\frac{T}{t_{FB}+t_{i}}*\frac{V*\alpha}{V_{FB}^{total}}}$= $\frac{n/V_{c}^{total}}{2*\frac{T}{t_{FB}+t_{i}}/V_{FB}^{total}}$

Notation and Definitions

*P*_v_： Production-efficiency-per-volume ratio between two-stage continuous and fed-batch fermentation;

T：Total biomanufacturing time;

t_0_：Fermentation time for initial-round cell growth when start the two-stage continuous fermentation;

t_c_：Cyclic time for the outflow of fermentation broth after saturated intracellular PHA accumulation within the two-stage continuous fermentation process;

$t_{FB}$：Single fed-batch fermentation time

t_i_：Interval time between two consecutive fed-batch fermentations (e.g., fermentation broth emptying, bioreactor cleaning, etc.)

V：Volume of the PHA-accumulation bioreactor for two-stage continuous fermentation, which is the same as the fed-batch one;

$V_{feed}$：Feeding volume in the PHA-accumulation bioreactor;

*α*：Fermentation broth loading coefficient (working volume to total volume)

*n*：Outflow frequency of two-stage continuous fermentation, calculated as:

n= $\frac{T-t_{0}}{t_{c}}$

$V_{c}^{total}$：Total bioreactor volume required for two-stage continuous fermentation, calculated as:

$$V_{c}^{total}=V+V_{c}^{cell grow}+V_{c}^{seed}$$

$V_{c}^{cell grow}$: Volume of the cell-growth bioreactor for two-stage continuous fermentation, calculated as:

$$V_{c}^{cell grow}=\frac{V*\alpha-V_{feed}}{\alpha}$$

$V_{c}^{seed}$: Volume of the seed-culture bioreactor for two-stage continuous fermentation, calculated as:

$$V_{c}^{seed}=\frac{V*\alpha-V_{feed}}{\alpha}*\frac{10\%}{n}$$

$V_{FB}^{total}$：Total bioreactor volume required for fed-batch fermentation, calculated as:

$V_{FB}^{total}=2V$+$V_{FB}^{seed}$

$V_{FB}^{seed}$: Volume of the seed-culture bioreactor for fed-batch fermentation, calculated as:

$V_{FB}^{seed}=2V$*10%

For two-stage continuous fermentation in this study, the fermentation time of cell-growth and PHA-accumulation state in two independent bioreactors was set to the same value (***t_0_ = t_c_***). For *P*_v_ ratio calculation, ***t_FB_*** and ***t_i_*** were set as 40 h and 8 h. The loading coefficient (α) for each bioreactor was 0.8. In the two-stage continuous system, the feeding volume (***V_feed_***) in the PHA-accumulation bioreactor was defined as one-fifth of the working volume, i.e., ***V_feed_ =*** $\frac{\boldsymbol{1}}{\boldsymbol{5}}$**V*α**. Under these assumptions, the function for *P*_v_ = *f*(T) can be simplified as follow:

*P*_v_$\boldsymbol{=}\frac{\frac{\boldsymbol{T-t}}{\boldsymbol{t}}\boldsymbol{/(1.8+}\frac{\boldsymbol{8\%}}{\frac{\boldsymbol{T-t}}{\boldsymbol{t}}}\boldsymbol{)}}{\frac{\boldsymbol{T}}{\boldsymbol{24}}\boldsymbol{/2.2}}$

The cyclic frequency of the outflow fermentation broth from PHA-accumulation bioreactor during the two-stage continuous fermentation typically ranges from 20 to 24 h. Calculated function curves were provided in Fig. 6e.

For comparative analysis between the experiment result against the function calculation, the experimental fitting analysis of *P*_v_ was obtained using the following equation:

*P_v_*=$\frac{V*\alpha\sum_{1}^{j} {PHA}_{i}/(1.8+\frac{0.08}{n})V}{V*\alpha*2*{PHA}_{FB}\frac{T}{40+8}/2.2V}$

where PHA_i_ represents the PHA titer for each cyclic outflow fermentation broth during the two-stage continuous fermentation, and ${PHA}_{FB}$ indicates the PHA titer for each fed-batch fermentation.

The equation can be thus simplified as follows:

*P*_v_ = $\frac{\sum_{1}^{j} {PHA}_{i}/(1.8+\frac{0.08}{n})}{{PHA}_{FB}\frac{T}{24}/2.2}$

### Appendix: Genes used in this study.

| **Genes** | **Descriptions** | **References** |
| --- | --- | --- |
| *dhaB-gdrAB* | Encoding glycerol dehydratase and its activator from *Klebsiella pneumoniae.* | ^11, 12, 13^ |
| *pduP_ST_* | Encoding coA-propanoylating propanal dehydrogenase from *Salmonella typhimurium.* | ^14, 15^ |
| *pduP_LR_* | Encoding coA-propanoylating propanal dehydrogenase from *Lactobacillus reuteri* | ^16^ |
| *xylA* | Encoding xylose isomerase from *Escherichia coli* MG1655. | ^17^ |
| *xfp* | Encoding phosphoketolase from *Lactobacillus rhamnosus* CGMCC1.120. | ^18^ |
| *aldD_Pp_* | Encoding aldehyde dehydrogenase from *Pseudomonas putida* KT2440, GenBank No.: AAN66172.1. | Lab stock |
| *ydcW_Kp_* | Encoding aldehyde dehydrogenase from *Klebsiella pneumoniae.* | ^13, 18^ |
| *aldD_Hb_* | Encoding aldehyde dehydrogenase from *Halomonas bluephagenesis*TD01. | ^19^ |
| *aldH_Ec_* | Encoding aldehyde dehydrogenase from *Escherichia coli* MG1655. | ^20^ |
| *gabD4_Re_* | Encoding aldehyde dehydrogenase from *Ralstonia eutropha* H16 with the mutant of E209Q and E269Q | ^21^ |
| *dhaT_Pp_* | Encoding 1,3-Propanediol dehydrogenase from *Pseudomonas. putida* KT2440, GenBank No.: AAN68411.1. | Lab stock |
| *adhP_Hb_* | Encoding alcohol dehydrogenase from *Halomonas bluephagenesis*TD01. | ^19^ |
| *4hbd* | Encoding 4-hydroxybutyrate dehydrogenase from *C. kluyveri*. | Lab stock |
| *sucD* | Encoding succinate semialdehyde dehydrogenase from *C. kluyveri Caulobacter.* | Lab stock |
| *ogdA* | Encoding 2-oxoglutarate decarboxylase from cyanobacterial, *Synechococcus sp.* PCC 7002. | Lab stock |
| *phaC_61-3_* | Encoding PHA synthase from *Pseudomonas entomophila* 61-3. | Lab stock |
| *phaC_Re_* | Encoding PHA synthase from *Ralstonia eutropha.* | Lab stock |
| *phaC_AR_* | Encoding a phaC chimeric enzyme consisting of 26% N-terminal sequence of *PhaC_Ac_* and 74% C-terminal sequence of *PhaC_Re_* derived from *Ralstonia eutropha.* | ^22^ |
| *phaC_Ah_* | Encoding PHA synthase from *Aeromonas hydrophila* 4AK4 | ^23^ |
| *PhaJ_Pe_* | Encoding enoyl-CoA hydratase from *Pseudomonas entomophila.* | Lab stock |
| *PhaJ_Ah_* | Encoding enoyl-CoA hydratase from *Aeromonas hydrophila* 4AK4. | ^22^ |

### Reference

1. Simon, R., Priefer, U. & Pühler, A. A broad host range mobilization system for in vivo genetic engineering: transposon mutagenesis in gram negative bacteria. *Bio/Technology* **1**, 784–791 (1983).
2. Tan, D., Xue, Y.-S., Aibaidula, G. & Chen, G.-Q. Unsterile and continuous production of polyhydroxybutyrate by *Halomonas* TD01. *Bioresour. Technol.* **102**, 8130–8136 (2011).
3. Shen, R., et al. Promoter engineering for enhanced P (3HB-*co*-4HB) production by *Halomonas* *bluephagenesis.* *ACS Synth. Biol.* **7**, 1897–1906 (2018).
4. Silva-Rocha, R., et al. The Standard European Vector Architecture (SEVA): a coherent platform for the analysis and deployment of complex prokaryotic phenotypes. *Nucleic Acids Res.* **41**, D666–D675 (2013).
5. Yin, J., Fu, X.-Z., Wu, Q., Chen, J.-C. & Chen, G.-Q. Development of an enhanced chromosomal expression system based on porin synthesis operon for halophile *Halomonas* sp. *Appl. Microbiol. Biotechnol.* **98**, 8987–8997 (2014).
6. Cai, L., et al. Comparative genomics study of polyhydroxyalkanoates (PHA) and ectoine relevant genes from *Halomonas* sp. TD01 revealed extensive horizontal gene transfer events and co-evolutionary relationships. *Microb. Cell Fact.* **10**, 88 (2011).
7. Tian, F., et al. Isolation, cloning and characterization of an azoreductase and the effect of salinity on its expression in a halophilic bacterium. *Int. J. Biol. Macromol.* **123**, 1062–1069 (2019).
8. Zhang, L., et al. Effective production of Poly (3-hydroxybutyrate-co-4-hydroxybutyrate) by engineered *Halomonas bluephagenesis* grown on glucose and 1, 4-Butanediol. *Bioresour. Technol.* **355**, 127270 (2022).
9. Reese, M. G. Application of a time-delay neural network to promoter annotation in the *Drosophila melanogaster* genome. *Comput. Chem.* **26**, 51–56 (2001).
10. Salgado, H., et al. RegulonDB v12. 0: a comprehensive resource of transcriptional regulation in *E. coli* K-12. *Nucleic Acids Res.* **52**, D255–D264 (2024).
11. Zhang, Y., et al. Sustainable biosynthesis of 3-hydroxypropionic acid from crude glycerol: Metabolic engineering and process optimization. *J. Clean. Prod.* **383**, 135524 (2023).
12. Qi, X., Guo, Q., Wei, Y., Xu, H. & Huang, R. Enhancement of pH stability and activity of glycerol dehydratase from *Klebsiella pneumoniae* by rational design. *Biotechnol. Lett.* **34**, 339–346 (2012).
13. Kim, J. W., Ko, Y. S., Chae, T. U. & Lee, S. Y. High‐level production of 3‐hydroxypropionic acid from glycerol as a sole carbon source using metabolically engineered *Escherichia coli.* *Biotechnol. Bioeng.* **117**, 2139–2152 (2020).
14. Wang, Q., et al. Biosynthesis of poly (3-hydroxypropionate) from glycerol by recombinant *Escherichia coli*. *Bioresour. Technol.* **131**, 548–551 (2013).
15. Meng, D.-C., et al. Production of poly (3-hydroxypropionate) and poly (3-hydroxybutyrate-co-3-hydroxypropionate) from glucose by engineering *Escherichia coli*. *Metab. Eng.* **29**, 189–195 (2015).
16. Luo, L. H., et al. Identification and characterization of the propanediol utilization protein PduP of *Lactobacillus reuteri* for 3-hydroxypropionic acid production from glycerol. *Appl. Microbiol. Biotechnol.* **89**, 697–703 (2011).
17. Tan, B., Zheng, Y., Yan, H., Liu, Y. & Li, Z.-J. Metabolic engineering of *Halomonas bluephagenesis* to metabolize xylose for poly-3-hydroxybutyrate production. *Biochem. Eng. J.***187**, 108623 (2022).
18. Zhao, P., Ma, C., Xu, L. & Tian, P. Exploiting tandem repetitive promoters for high-level production of 3-hydroxypropionic acid. *Appl. Microbiol. Biotechnol.* **103**, 4017–4031 (2019).
19. Jiang, X.-R., Yan, X., Yu, L.-P., Liu, X.-Y. & Chen, G.-Q. Hyperproduction of 3-hydroxypropionate by *Halomonas bluephagenesis*. *Nat. Commun.* **12**, 1513 (2021).
20. Jo, J.-E., et al. Cloning, expression, and characterization of an aldehyde dehydrogenase from *Escherichia coli* K-12 that utilizes 3-hydroxypropionaldehyde as a substrate. *Appl. Microbiol. Biotechnol.* **81**, 51–60 (2008).
21. Honjo, H., Tsuruno, K., Tatsuke, T., Sato, M. & Hanai, T. Dual synthetic pathway for 3-hydroxypropionic acid production in engineered *Escherichia coli*. *J. Biosci. Bioeng.* **120**, 199–204 (2015).
22. Matsumoto, K. i., Takase, K., Yamamoto, Y., Doi, Y. & Taguchi, S. Chimeric enzyme composed of polyhydroxyalkanoate (PHA) synthases from *Ralstonia eutropha* and *Aeromonas caviae* enhances production of PHAs in recombinant *Escherichia coli*. *Biomacromolecules* **10**, 682–685 (2009).
23. Liu, F., et al. Metabolic engineering of *Aeromonas hydrophila* 4AK4 for production of copolymers of 3-hydroxybutyrate and medium-chain-length 3-hydroxyalkanoate. *Bioresour. Technol.* **102**, 8123–8129 (2011).
